# Gene turnover and contingency facilitated the repeated evolution of C_4_ photosynthesis in grasses

**DOI:** 10.1101/2025.04.23.650007

**Authors:** Lara Pereira, Ahmed S Alenazi, Sahr Mian, Ilia J Leitch, Pascal-Antoine Christin, Colin P Osborne, Luke T Dunning

## Abstract

**Summary:**

- In grasses, almost all species belong to two evenly sized clades (BOP and PACMAD), yet the >20 independent origins of C_4_ photosynthesis in this family only occur in the PACMAD lineage. Here, we identify potential genetic precursors for C_4_ photosynthesis that were present at the base of the PACMAD clade, representing the last common ancestor of all C_4_ grasses.
- We generated the first reference genomes for Aristidoideae species *(Aristida adscensionis* and *Stipagrostis hirtigluma)*, the sister lineage of all other PACMAD grasses. In combination with 34 other Poales genomes, we identify gene gains at the base of the PACMAD clade, and genes lost in the sister BOP lineage.
- Candidate C_4_ precursors include β-carbonic anhydrase, as well as genes involved in amino acid and nitrate transport, carbon metabolism, oxidative stress management and transcription regulation.
- Gene turnover created the necessary genetic contingency, facilitating the independent recruitment and refinement of C_4_ genes in multiple independent origins. Our results support the idea that the repeated evolution of C_4_ photosynthesis in PACMAD grasses was not driven by a single genetic event but was instead underwritten by chance genetic changes that originated long before there was selection for the trait itself.

## Introduction

Complex traits are composed of multiple components working in synergy to increase an organism’s fitness. By comparing different species, we can observe how these composite traits manifest into impressive phenotypic innovations and novel adaptations. However, the evolutionary dynamics behind the progressive emergence of complex traits are often unknown. The evolution of complex traits is typically envisaged as a stepwise progression of multiple intermediate stages, each offering an incremental fitness advantage that is selected for through a process of descent with modification (Darwin, 1859). The order that the individual components are assembled is of great interest (Meléndez-Hevia et al., 1996; Lenski et al., 2003), especially when a key innovation is foundational to the emergence of a particular trait (e.g. Ourisson and Nakatani, 1994). The presence of such evolutionary precursors may be one explanation for the non-random phylogenetic clustering of certain convergent traits (Christin et al., 2015).

Gene duplication is a major driving force in evolution because it generates novel genetic variation that selection can act upon to fuel functional innovation (Conant & Wolfe, 2008; Magadum et al., 2013). The loss of a gene duplicate is the most common outcome, but natural selection may act to preserve a duplicate if there is either: (i) a beneficial gene dosage effect, (ii) subfunctionalization (duplicate pairs each perform a subset of the original functions) or (iii) neofunctionalization (one duplicate acquires a novel function) (Monson, 1999; Conant & Wolfe, 2008; Panchy et al., 2016). All three mechanisms are involved in the process of generating novelty (Conant & Wolfe, 2008), although neofunctionalization is generally thought to be the most important for the emergence of novel complex traits (e.g. Chau et al., 2017; Huu et al., 2020). Gene duplications can originate through whole genome duplication (polyploidization) or small-scale duplications, typically either through unequal crossing-over events or mediated by transposable elements. Gene duplications have contributed to the developmental programme of many organisms, including the HOX genes responsible for mammalian body form (Wagner et al., 2003), and the developmental and regulatory loci underpinning the diversity of flowering plants (Moore and Purugganan, 2005).

In addition to gene duplication, new genes can evolve from noncoding DNA. This *de novo* gene birth can also generate the substrate for evolutionary innovation (Xia et al., 2025). For example, orphan genes in *Arabidopsis* and *Brassica rapa* have had an adaptive impact on starch (Li et al., 2009) and soluble sugar metabolism (Jiang et al., 2020), respectively. Pseudogenization or gene loss can also play a significant role in a species evolutionary trajectory. This could be through the loss of a genetic precursor or key part of a particular pathway, for example humans and some other mammals have convergently lost the ability to synthesise ascorbic acid due to a loss of function mutations in the *GLO* gene (Drouin et al., 2011). Conversely, gene loss can open new evolutionary paths to increased fitness by removing certain constraints (Helsen et al, 2020).

The C_4_ photosynthetic pathway is a complex trait that increases the efficiency of carbon fixation in hot and high light environments when compared to the ancestral C_3_ pathway (Ehleringer & Bjorkman, 1977; Ehleringer, 1978; Ehleringer et al., 1991, 1997; Sage et al., 2012; Atkinson et al., 2016). C_4_ photosynthesis requires the coordinated action of numerous anatomical and biochemical components to concentrate CO_2_ around the carbon-fixing enzyme Rubisco (Hatch, 1987). Gene duplication and subsequent neofunctionalization has previously been proposed as a mechanism for creating C_4_ expression patterns (Monson, 2003). However, there is a lack of evidence for the immediate role of gene duplication in C_4_ evolution (Gowik and Westhoff 2011; Williams et al. 2012; van den Bergh et al. 2014), with the recruitment of core metabolic genes based on their ancestral expression patterns and catalytic properties (Wang et al., 2009; Hibberd and Covshoff, 2010; Aubry et al., 2011; Christin et al., 2013a, 2015; Emms et al., 2016; Moreno-Villena et al., 2018; Dunning et al., 2019). Beyond the recruitment of core C_4_ enzymes, whole genome duplication is thought to have enabled the evolution of C_4_ anatomy and biochemistry in the eudicot *Gynandropsis gynandra* (Hoang et al., 2021), and a single photorespiratory gene duplication underpins the development of the photorespiratory CO_2_ pump in the C_3_-C_4_ intermediate eudicot *Flaveria* species (Schulze et al., 2013). In the grass *Alloteropsis semialata*, post co-option duplication and increased dosage seems to have played a role in reinforcing an initially weak C_4_ cycle (Bianconi et al., 2018). However, distinguishing whether genetic changes enabled the emergence of the C_4_ pathway, or occurred as a consequence of its establishment, remains a major challenge (Dunning et al., 2017).

Despite its apparent complexity, C_4_ photosynthesis has evolved over 60 times in angiosperms, representing a remarkable example of convergent evolution (Heyduk et al., 2019). The origins of C_4_ photosynthesis are phylogenetically clustered (Sage et al., 2011), with more than 20 C_4_ origins in grasses alone (Grass Phylogeny Working Group II, 2012). Even within the grass family (Poaceae), the multiple origins of C_4_ photosynthesis are clustered, only occurring in the **PACMAD** clade (six subfamilies: **P**anicoideae, **A**ristidoideae, **C**hloridoideae, **M**icrairoideae, **A**rundinoideae, and **D**anthonioideae) which is sister to the **BOP** clade (three subfamilies: **B**ambusoideae, **O**ryzoideae, and **P**ooideae) where C_4_ species are conspicuously absent (Grass Phylogeny Working Group II, 2012; Fig. 1). These two clades contain over 99% of described grass species and are equally sized (BOP = 5,941 species; PACMAD = 5,815 species) (Soreng et al., 2022).

**Figure 1.**
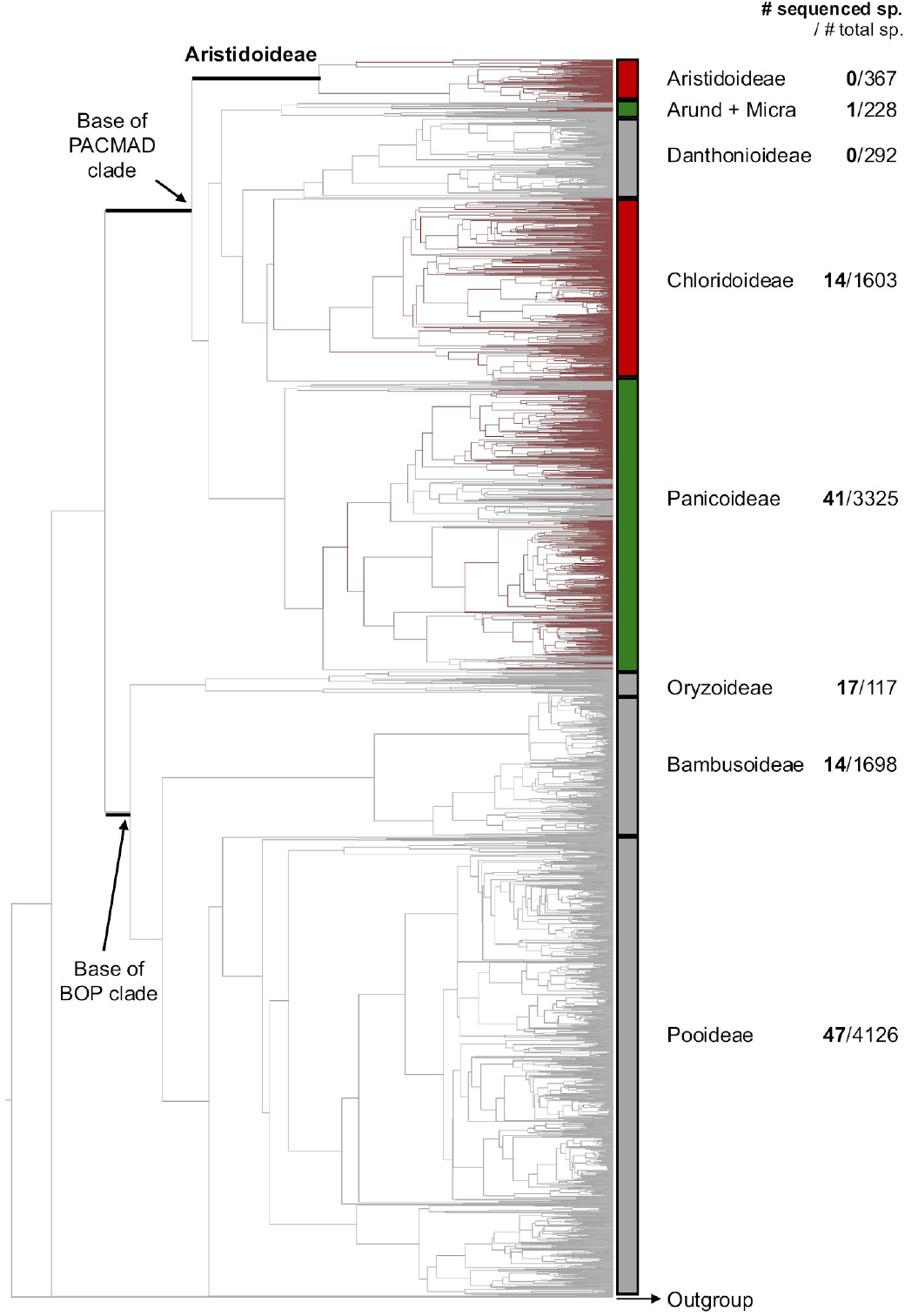
Phylogeny, C_4_ origins and sequenced genomes. The dated phylogeny is from Spriggs et al. (2014), with C_3_ species represented in grey, and C_4_ species, in red. Bars for each clade indicate if they are exclusively C_3_ (grey), C_4_ (red) or mixed (green). The number of species per subfamily was extracted from Soreng et al. (2022) and the number of sequenced genomes per subfamily was searched in NCBI (15/11/2024). Arund = Arundinoideae, Micra = Micrairoideae.

The absence of C_4_ photosynthesis in BOP grasses suggests that species from the PACMAD clade carry precursors that make them prone to evolve C_4_ (Fig. 1), driven by either gene gains specific to the PACMAD clade or gene losses in the BOP lineage. A previous phylogenomic study using transcriptome data found no evidence for a whole genome duplication event at the base of the PACMAD clade (Zhang et al., 2024), indicating that small-scale duplications are more likely to be the source of these precursors. Relevant precursors could be associated with the biochemical pathway itself, or related with leaf anatomy differences (e.g. proportion of the vascular bundle sheath layer) known to exist between the BOP and PACMAD clades (Christin et al., 2013b, 2015). The genetic control of leaf development has only been partially elucidated (Lewis and Hake, 2016, Sedelnikova et al., 2018), preventing a comprehensive inspection of specific genes that might be responsible for these preadaptations. A global analysis using comparative genomics to identify gene families that have been duplicated and retained in two independent C_4_ origins in a single grass subfamily (Panicoideae) unveiled 41 orthologous genes with heterogeneous functions, including transcription factors, transporters, and genes involved in hormone metabolism (Emms et al., 2016), with these genes potentially acting as genetic precursors to C_4_ evolution. Whilst there are high-quality reference genomes for more than one hundred grass species (Fig. 1), they are only available for one of the two groups resulting from the earliest split within PACMAD lineage. Genomes for representatives of the Aristidoideae subfamily are lacking, preventing a global analysis of the specific precursors present in the last common ancestor of all C_4_ origins in grasses.

In this study, we aimed to identify gene gains or losses that may represent preadaptations to C_4_ photosynthesis present in the last common ancestor of the PACMAD clade. To achieve this, we generated reference genomes for two members of the Aristidoideae family, *Aristida adscensionis* and *Stipagrostis hirtigluma*. We then used these genomes, together with publicly available assemblies for other grass species, to identify duplication and *de novo* gene births at the base of the PACMAD clade, and gene losses at the base of the BOP lineage. We found that the duplicated genes were enriched for transmembrane transporters and genes associated with resistance to oxidative damage, functions that may be foundational to the evolvability of C_4_ photosynthesis in the PACMAD grasses.

## Materials and methods

### Tissue collection and DNA extractions

*Aristida adscensionis* seeds were obtained from the USDA Germplasm Resources Information Network (GRIN) (accession number = PI 269867, collection location = Pakistan), and *Stipagrostis hirtigluma* var. *patula* seeds from the Royal Botanic Gardens, Kew Millenium Seed Bank (accession number = 74902, collection location = Botswana). Seeds were germinated and grown in a glasshouse at the Arthur Willis Environment Centre, University of Sheffield (UK) under semi-controlled conditions (12 h daylight, 25/20 °C day/night temperature). The 1C genome size for each species was estimated by flow cytometry using the one-step protocol (Doležel et al., 2007) with minor modifications (see Clark et al., 2016). *Petroselinum crispum* ‘Champion Moss Curled’ (2C = 4.50 pg; Obermayer et al., 2002) was used as the calibration standard and nuclei were isolated in the General Purpose Buffer (GPB; Loureiro et al. 2007), supplemented with 3% PVP and 0.08% (v/v) beta-mercaptoethanol.

High molecular weight genomic DNA was extracted for each species from flash frozen leaf tissue stored at -80 °C using the NucleoBond® HMW DNA kit (Macherey-Nagel), following the manufacturer’s protocol. DNA extractions were purified using AMPure XP Beads (Beckman Coulter Life Sciences). DNA length, purity and concentration were estimated with a Femto Pulse (Agilent), NanoDrop (Thermo Fisher Scientific), and Qubit (Thermo Fisher Scientific), respectively. PacBio HiFi DNA libraries were prepared and run on Sequel II SMRT Cells in CCS mode at the NERC Centre for Genomic Research (University of Liverpool, UK). In total, we ran two SMRT cells for *A. adscensionis* and three SMRT cells for *S. hirtigluma*.

#### Organelle assembly

To remove organelle data prior to nuclear genome assembly, we first generated reference plastid and mitochondrial organelle genomes. Plastid sequences were identified by aligning the raw HiFi reads with BLASR v5.3.5-0 (Bussotti et al., 2011) to published plastome assemblies for each species downloaded from NCBI GenBank *(A. adscensionis =* MZ373986.1; *S. pennata =* MZ373985.1). To be considered for downstream analysis, reads were filtered by alignment length (≥5 kb) and alignment similarity (≥99%). These reads were then *de novo* assembled using hifiasm vθ.16.1 (Cheng et al., 2021) with default parameters. Primary contigs were inspected using circlator clean vl.5.5 (Hunt et al., 2015), discarding shorter contigs that were nested within larger sequences. The remaining contigs were annotated with the GeSeq v2.03 (Tillich et al., 2017) web portal. The annotations were used to identify overlapping contigs that were then manually trimmed, merged and circularised in Geneious Prime (v2023.0.1, Build 2022-11-28) to obtain the complete plastid genome, before being re-annotated with GeSeq.

Assembling mitochondria can be problematic for plants because they are not necessarily circular and can exist in multiple conformations within a cell (Sloan et al., 2012). To assemble mitochondria, we used MitoHiFi v3.2 (Uliano-Silva et al., 2023) with a minimum percentage of each HiFi read sequence in the blast match to a closely related mitochondrial sequence (-p) set to 30%. As expected, a single fully resolved mitochondrial genome was not obtained. To select unique, long contigs, we used circlator clean and annotated the remaining contigs with GeSeq. Only contigs containing at least one gene model were included in the draft mitochondrial contigs.

### Nuclear assembly

The estimated coverage and ploidy for each species were calculated using the k-mer spectrum generated with GenomeScope 2.0 (Ranallo-Benavidez et al., 2020). Prior to nuclear genome assembly, we removed reads belonging to the organelles by mapping raw HiFi reads to the plastid and mitochondrial genomes with minimap2 v2-2.24 (Li, 2018), discarding those where ≥80% of the read aligned. The organelle-filtered HiFi reads were then *de novo* assembled using hifiasm, using default parameters except for the number of haplotypes in *S. hirtigluma* being set to four as it was inferred to be tetraploid (see results below). The assembly completeness was checked using BUSCO v5.4.2 (Simão et al., 2015) with the poalesodblO database, and Inspector vl.2 (Chen et al., 2021) was used to estimate assembly quality values (QV).

### Genome annotation

Extensive *de-novo* TE Annotator vl.9.4 (EDTA) (Ou et al., 2019) was used to annotate transposable elements (TEs), and RepeatMasker v4.1.2-pl to annotate simple tandem repeats and low complexity (polypurine, AT-rich) regions, with default parameters. The identified TEs and simple repeats were then soft-masked in each draft genome using BEDtools maskfasta (Quinlan and Hall, 2010). To annotate the protein coding genes in the soft-masked reference genome, we used Helixer v.0.3.2, a tool that uses a deep learning model to annotate large eukaryotic genomes (Stiehler et al., 2020, Holst et al., 2023).

### Identifying orthologous genes and duplication events

We used OrthoFinder v.2.5.4 (Emms and Kelly, 2019) to identify groups of orthologs and duplication events in grasses. We included the two reference genomes produced here in addition to 34 published genomes from across the Poales (15 x C_4_ PACMAD, 4 x C_3_ PACMAD, 9 x BOP, 2 x early diverging grasses and 4 x non-grass Poales) with the aim of representing as many subfamilies and C_4_ origins as possible (Fig. 2; Table SI). To reduce the effect of annotation quality on our interpretations we re-annotated the published genomes with Helixer when the extracted coding sequences (CDS) were < 90% complete according to BUSCO analysis (Table S2). Poaceae genomes had to have ≥90% complete BUSCO annotation scores to be included in the final analysis (Table S2), an exception was made for the *Alloteropsis semialata* genome (>89%) as there are relatively few C_3_ PACMAD species with genomes available and this species is a model for C_4_ evolution in grasses (Pereira et al., 2023). For Poales outgroups, the completeness threshold was lowered *(Carex scoparia* [Cyperaceae] 79.8% complete BUSCO; *Juncus effusus* [Juncaceae] 78.1% complete BUSCO) as it has been previously suggested that their relatively low BUSCO scores are a result of only Poaceae and Bromeliaceae species being used in the construction of the Poales BUSCO gene database, meaning these genes are not necessarily conserved across the whole order (Planta et al., 2022).

**Figure 2.**
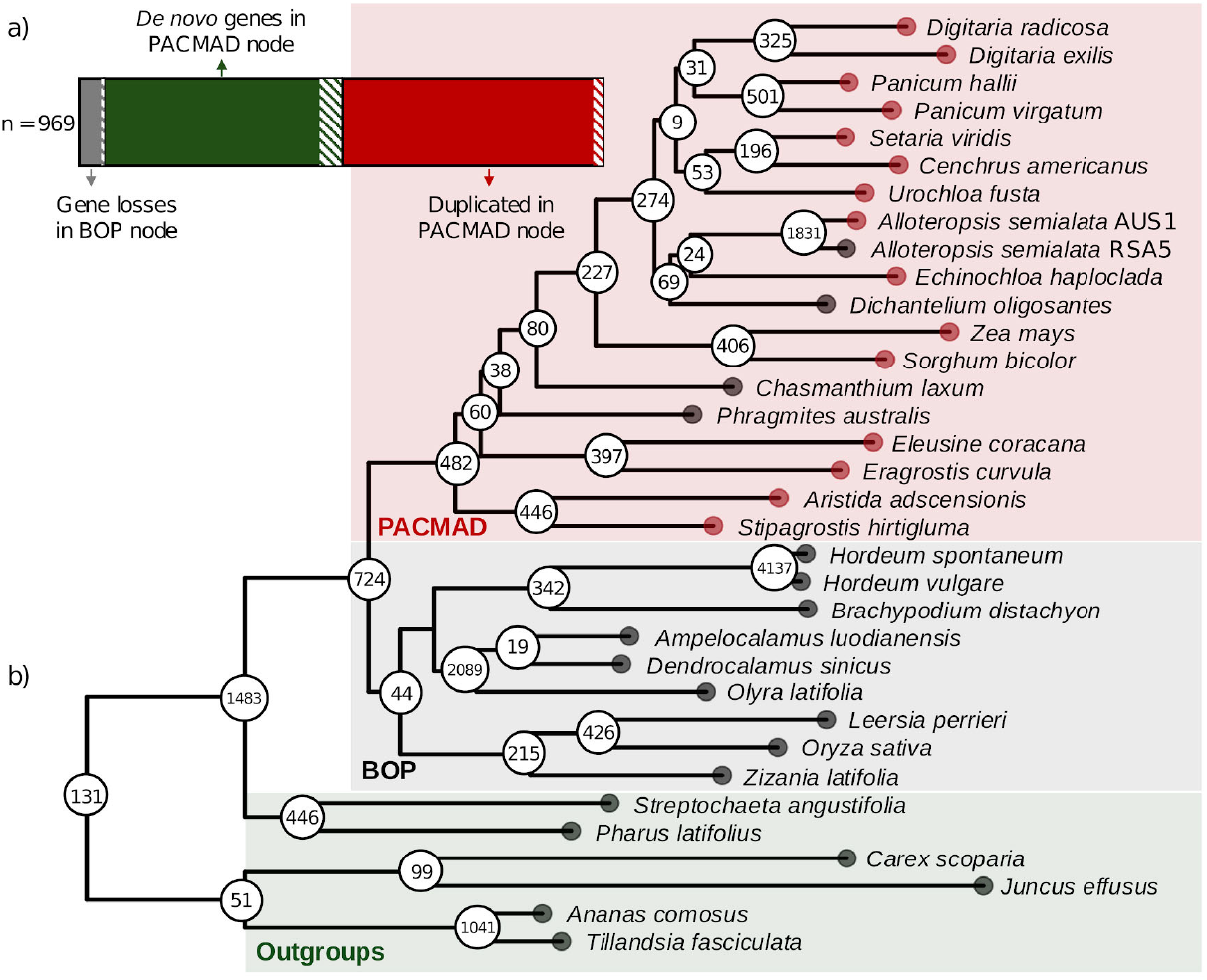
Candidate genetic precursors of C_4_ photosynthesis in grasses, (a) The proportion of candidate genes arising through gene duplication, gene loss and *de novo* origin. The dashed portion of each group represents the core set for each group, i.e. present in all 15 C_4_ PACMAD species used in this study, (b) Species tree and number of duplications per node generated by OrthoFinder. C_4_ species represented by a red circle at the tip, and C_3_ by a black circle.

Orthologs across all 36 genomes were identified with OrthoFinder using the CDS, aligned using MUSCLE v3.2 (Edgar, 2004), and maximum likelihood trees inferred using FastTree v2 (Price et al., 2010). OrthoFinder generates a species tree inferred from all orthogroups using the ‘Species Tree Inference from All Genes’ (STAG) method (Emms and Kelly, 2018), rooted using ‘Species Tree Root Inference from Gene Duplication Events’ (STRIDE) (Emms and Kelly 2017). OrthoFinder then reconciles gene duplications against the inferred STAG species tree to determine the branch on which the duplication occurred, with a minimum support value of ≥50% (i.e. ≥50% of species that share a common ancestor on that branch have retained both copies of the duplicated gene). Of particular interest for this study are the orthogroups inferred to have been duplicated at the base of the PACMAD clade, the branch representing the last common ancestor of all grass C_4_ origins. To determine if the number of duplications along the PACMAD branch was higher than expected, we calculated the rate of duplication across the STAG tree using divergence times for homologous branches extracted from a previously dated phylogeny (Christin et al., 2014).

The PACMAD specific duplicates were further filtered to a core set, i.e. they had to be [1] present in all 15 C_4_ species included in the analysis, and [2] verified in phylogenetic trees constructed independently of OrthoFinder. To infer individual gene trees, we first identified homologous sequences in the 36 species with BLASTn v2.16.0 using all coding sequences from the orthogroup of interest as a query. Blast matches had to be ≥300 bp long, and we only retained the longest match for each sequence in the database. The BLASTn matching regions were then aligned with mafft v7.525 (Katoh & Standley, 2013), set to automatically choose the best alignment algorithm (--auto). A phylogenetic tree was then inferred with PhyML v.21031022 (Guindon et al., 2010), with the best substitution model identified using Smart Model Selection (SMS) v.1.8.1 (Lefort et al., 2017).

Finally, for the verified set of PACMAD duplicates, we tested if this resulted in a significantly increased gene copy number at the orthogroup level. For each orthogroup the gene copy number was normalised by ploidy level, and it was determined if the data was normally distributed using the Shapiro-Wilk test (p < 0.05). Differences in gene copy number between BOP and PACMAD species were assessed using two-tailed t tests (normal data) or Wilcoxon signed-rank test (non-normal data), with Benjamini-Hochberg correction of multiple testing (significance at adjusted p < 0.05). All analyses were done in R.

### *Gene losses in the BOP clade and* de novo *genes in the PACMAD clade*

Orthgroups lost in the BOP clade were identified as those present in at least 1 of the 6 outgroup species and in ≥50% of PACMAD species (including at least 1 representative from each of the Panicoideae, Aristidoideae, and Chloridoideae subfamilies), but absent in all 9 BOP species analysed. PACMAD-specific *de novo* orthogroups were identified as those absent in the outgroup and BOP species, and present in ≥50% of PACMAD species (including at least 1 representative from each of the Panicoideae, Aristidoideae, and Chloridoideae subfamilies). Both orthogroups lost in the BOP clade and PACMAD-specific orthogroups were further filtered to a core set (i.e. present in all 15 C_4_ species included in the analysis).

### Gene ontology

To determine if certain Gene Ontology (GO) terms (Ashbumer et al., 2000) are overrepresented in the genes that are either duplicated at the base of the PACMAD clade, lost in the BOP clade, or are PACMAD-specific *de novo* genes, we used the statistical overrepresentation test in the Panther online tool v.19 (https://pantherdb.org/) (Thomas et al., 2021; Mi et al., 2013). Maize was used as the background reference genome for the analysis as it is the best characterised PACMAD species, and the target lists used either [1] all maize loci in the orthogroups of interest (BOP losses and *de novo* PACMAD), or [2] one randomly selected duplicate from each of the PACMAD specific duplication events. A Fisher’s exact test was used to determine which PANTHER GO-Slim terms were significantly enriched (FDR < 0.05) in the target lists. The results were further restricted to only include those annotated with 10 to 500 genes in the maize genome, this excluded overly general categories and those with low statistical support. Finally, QuickGO (https://www.ebi.ac.uk/QuickGO/) was used to generate a hierarchical representation of the significant GO terms.

The function of the candidate C_4_ precursors (i.e. PACMAD-specific duplicates, PACMAD de novo genes, and BOP losses) were further investigated using BLASTp v2.16.0 against the SwissProt database. A randomly selected coding sequence from each orthogroup (or duplicate) was translated with EMBOSS Transeq v.6.6.0 (Rice et al., 2000) and used as a protein query, with -maxtargetseqs set to 3 and an E-value cut-off of lxlO”^5^. We further validated the function of the core sets of PACMAD duplicates, BOP losses and *de novo* PACMAD genes using Ensembl Plants (Howe et al., 2020), the maize database MaizeGBD (Woodhouse et al., 2021), the *Arabidopsis thaliana* database TAIR (Berardini et al., 2015), and literature searches.

### *Tissue-specific C*_*4*_ *expression patterns*

The C_4_ cycle involves the recruitment and compartmentalisation of multiple enzymes across mesophyll (M) and bundle sheath (BS) tissues. Gene duplication may facilitate the required tissue-specific subfunctionalization, enabling loci to separate their role in C_4_ photosynthesis from other functions that they may perform. We used a chi-squared test to determine if genes resulting from PACMAD-specific duplications are more likely to be differentially expressed between M and BS cells than the genome-wide proportion. To do this we cross-referenced our candidate gene list with the results from previous studies in maize (Li et al., 2010) and sorghum (Döring et al., 2016) that identified genes significantly differentially expressed between M and BS tissues. If the gene models for our candidate loci had changed as a result of updated genome annotations, we applied the same expression pattern to the fused or split gene models used in our analysis. We also included any genes arising from further duplications of these loci that may have occurred after PACMAD lineages began diverging. To infer whether some of these candidates changed their pattern of expression after duplication, we also used an available transcriptomics dataset in rice (Hua et al., 2021).

## Results

### Reference genomes for two Aristidoideae grasses

We assembled reference genomes for two divergent Aristidoideae species, *Aristida adscensionis* and *Stipagrostis hirtiglumă*, representing independent realisations of the C_4_ phenotype. For *A. adscensionis*, we generated 31.0 Gb of HiFi data, which is approximately 53x coverage of the nuclear genome based on the flow cytometry estimate for 1C of 0.58 Gb (Table 1). The analysis of the k-mer spectrum suggested that the *A. adscensionis* individual sequenced was diploid (Fig. SI), and the final assembly was 0.45 Gb (77.8% of the genome size estimated using flow cytometry), relatively contiguous (186 contigs andN50 = 5.6 Mb) and 98.1% of BUSCOs were complete (Table 1). Genome annotation showed that 53.0% of the genome is made up of TEs, 0.81% simple tandem repeats, 0.08% low complexity regions, and we were able to identify 32,006 protein coding genes (Table 1).

**Table 1.** Assembly statistics for the two Aristidoideae genomes.

|  |  | *Aristida*<br>*adscensionis* | *Stipagrostis*<br>*hirtigluma* |
| --- | --- | --- | --- |
| Sequencing | Number of reads | 1,889,280 | 4,507,740 |
|  | Average read length | 16.39 kb | 15.01 kb |
|  | Sequence data volume | 30.97 Gb | 67.68 Gb |
| Assembly | Ploidy | Diploid | Tetraploid |
|  | Estimated 1C genome size | 0.58 Gb | 0.48 Gb* |
|  | Total assembly length | 0.45 Gb | 0.61 Gb |
|  | Number of contigs | 186 | 458 |
|  | Contig N50 | 5.6 Mb | 12.14 Mb |
|  | Longest contig | 17.85 Mb | 31 Mb |
|  | Assembly quality value | 63.55 | 25.06 |
|  | BUSCO Complete (Duplicated) | 98.1% (2.4%) | 98.7% (35.1%) |
|  | BUSCO Fragmented | 0.9% | 0.8% |
| Annotation | Transposable Elements | 53.0% | 48.7% |
|  | Simple tandem repeats | 0.81% | 0.72% |
|  | Low complexity regions | 0.08% | 0.07% |
|  | Number of gene models | 32,006 | 50,517 |

\*Monoploid genome size (1Cx)

We generated 67.7 Gb of HiFi data for *S. hirtiglumă* (Table 1). The analysis of the k-mer spectrum suggested it was a tetraploid (unknown if alio- or auto-polyploid) (Fig. S2), therefore the monoploid genome size (i.e. the lCx value corresponding to the amount of DNA in one chromosome set calculated from the genome size measured by flow cytometry) was estimated to be 0.48 Gb (corresponding to 141x coverage). The final assembly was 0.61 Gb, which is higher than the monoploid flow cytometry estimate, possibly because divergent sequences from different subgenomes are assembled separately and concatenated in the final assembly. The assembly is more fragmented (458 contigs, N50 = 12.1 Mb) than the *A. adscensionis* assembly, despite the higher coverage and larger N50 obtained. This is most likely due to the added complexity of assembling a polyploid species. In terms of completeness, 98.7% of BUSCOs were complete (Table 1), and 35.1% were duplicated, further supporting the partial redundancy of the assembly. Genome annotation showed that 48.72% of the genome is made up of TEs, 0.72% simple tandem repeats, 0.07% low complexity regions, and we were able to identify 50,517 protein coding genes (Table 1).

### Identification of genetic precursors of C_4_ photosynthesis in grasses

In total we identified 969 candidate C_4_ genetic precursors (Fig. 2a), including 482 genes duplicated in the last common ancestor of PACMAD (Table S3), 440 orthogroups that formed *de novo* in the last common ancestor of PACMAD (Table S4), and 47 orthogroups that were lost in the last common ancestor of BOP (Table S5). The number of predicted duplications varied across the phylogeny (Fig. 2b; Table S6), with a mean rate of duplication for each branch being 24.7 duplications per Ma (SD = 18.4). The branch leading to the PACMAD clade does not have a significantly higher duplication rate than the rest of the phylogeny (38.8 duplications per Ma; 79^th^ percentile), ruling out a burst of gene duplication at the base of the PACMAD clade.

Significant enrichment of GO terms was only detected for the genes duplicated at the base of the PACMAD clade. The 23 significantly enriched GO terms were associated with multiple transporter and enzymatic activities that are essential for metabolism, detoxification, and cellular homeostasis (Table 2, Fig. S3). Of particular relevance to the evolution of C_4_ photosynthesis is the enrichment of GO terms associated with carboxylic acid (G0:0046943, G0:0046942) and amino acid (G0:0015171, G0:0015179) transmembrane transport, oxidoreductase activity (G0:0016614, G0:0016616, G0:0006749), and active transmembrane transport against concentration gradients (G0:0042626, G0:0015399).

**Table 2.** GO terms significantly overrepresented in the PACMAD specific gene duplicates.

| GO Term | Description | Class | #Ref | #Dup | FE | Padj |
| --- | --- | --- | --- | --- | --- | --- |
| *GO terms associated with transporter activity* |  |  |  |  |  |  |
| GO:0008028 | Monocarboxylic acid transmembrane transporter activity | MF | 18 | 3 | 16.46 | 1.47E-02 |
| GO:0015179 | L-amino acid transmembrane transporter activity | MF | 22 | 3 | 13.47 | 2.49E-02 |
| GO:0015171 | Amino acid transmembrane transporter activity | MF | 89 | 7 | 7.77 | 1.42E-03 |
| GO:0019829 | ATPase-coupled monoatomic cation transmembrane transporter activity | MF | 54 | 4 | 7.32 | 3.70E-02 |
| GO:0046943 | Carboxylic acid transmembrane transporter activity | MF | 109 | 7 | 6.34 | 3.27E-03 |
| GO:0005342 | Organic acid transmembrane transporter activity | MF | 109 | 7 | 6.34 | 3.11E-03 |
| GO:0042626 | ATPase-coupled transmembrane transporter activity | MF | 133 | 7 | 5.20 | 9.58E-03 |
| GO:0008514 | Organic anion transmembrane transporter activity | MF | 142 | 7 | 4.87 | 1.30E-02 |
| GO:0015399 | Primary active transmembrane transporter activity | MF | 151 | 7 | 4.58 | 1.72E-02 |
| GO:0006865 | Amino acid transport | BP | 86 | 7 | 8.04 | 3.84E-02 |
| GO:0046942 | Carboxylic acid transport | BP | 107 | 7 | 6.46 | 3.89E-02 |
| GO:0015849 | Organic acid transport | BP | 107 | 7 | 6.46 | 3.11E-02 |
| *GO terms associated with catalytic activity (including response to oxidative stress)* |  |  |  |  |  |  |
| GO:0004364 | Glutathione transferase activity | MF | 72 | 6 | 8.23 | 2.50E-03 |
| GO:0035251 | UDP-glucosyltransferase activity | MF | 121 | 9 | 7.35 | 2.08E-04 |
| GO:0016616 | Oxidoreductase activity, acting on the CH-OH group of donors, NAD or NADP as acceptor | MF | 192 | 12 | 6.17 | 6.91E-05 |
| GO:0008194 | UDP-glycosyltransferase activity | MF | 149 | 9 | 5.97 | 1.02E-03 |
| GO:0016614 | Oxidoreductase activity, acting on CH-OH group of donors | MF | 199 | 12 | 5.96 | 7.21E-05 |
| GO:0046527 | Glucosyltransferase activity | MF | 158 | 9 | 5.63 | 1.05E-03 |
| GO:0016765 | Transferase activity, transferring alkyl or aryl (other than methyl) groups | MF | 151 | 8 | 5.23 | 3.78E-03 |
| GO:0016758 | Hexosyltransferase activity | MF | 274 | 10 | 3.61 | 1.13E-02 |
| GO:0016757 | Glycosyltransferase activity | MF | 381 | 11 | 2.85 | 3.34E-02 |
| GO:0006749 | Glutathione metabolic process | BP | 66 | 6 | 8.98 | 3.88E-02 |
| GO:0006790 | Sulfur compound metabolic process | BP | 180 | 9 | 4.94 | 4.56E-02 |

#Ref = number of genes in reference list; #Dup = number of duplicated genes; FE = fold enrichment compared to the expected value; Padj = Benjamini-Hochberg adjusted p-value.

### Core C_4_ photosynthesis genetic precursors

Among the candidate genetic precursors for C_4_ photosynthesis we identified (PACMAD duplicates, PACMAD *de novo* genes, BOP losses), those consistently retained across all C_4_ species are more likely to have contributed to the repeated evolution of C_4_ photosynthesis in grasses. In total, both copies resulting from 3.7% (n = 18) of PACMAD gene duplications, 9.5% (n = 42) of PACMAD *de novo* orthogroups, and 12.7% (n = 6) of BOP orthogroup losses were retained in all 15 C_4_ species used in this study (Table S8).

The core set of 18 PACMAD duplicated and C_4_ retained candidates was further validated using additional gene trees (Fig. S4 & Dataset SI), with 15 duplication events validated and 3 unconfirmed: OG0003655 contained too few sequences (n = 26), all from PACMAD species; OG0001524 topology was inconsistent with the species tree, with very low support; and OG0002361 duplication possibly occurred prior to the BOP and PACMAD clades diverging (Dataset SI). Finally, we looked at the overall gene copy number in the orthogroups that contained the 15 core PACMAD-specific duplication events. There was a lot of variation between species which presumably results from other independent duplications and/or losses (Fig. S5). However, there was a significantly higher (adjusted p-value <0.05) gene copy number in PACMAD compared to BOP in 11 of these orthogroups (Table S7, Fig. S5).

A majority of the 15 core PACMAD duplicated genes are involved with transport, carbon metabolism or reducing the effects of oxidative damage (Fig. 3 and Table S8). The five transporters move amino acids (OG0002111 & OG0000861), nitrates (OG0000391 & OG0000379), and toxic compounds (OG0000180). The three carbon metabolism enzymes are: phosphoglycerate mutase-like (PGML) protein (OG0001190) - glycolysis/gluconeogenesis & Calvin-Benson-Bassham cycle; (DL)-glycerol-3-phosphatase (OG0003194) - glycolysis & potentially photorespiration; and a chloroplastic aspartyl proteinase NANA protein (OG0000505), that potentially plays a role in photoassimilate partitioning and the cross talk between the chloroplast and nucleus (Paparelli et al., 2012). The three enzymes with functions relating to reducing oxidative damage are: a peroxidase (OG0000809) linked to lignin biosynthesis (Femández-Pérez et al., 2015); a salutaridine reductase (SALR) (OGOOOOIOI) that is orthologous to an Arabidopsis NAD(P)-binding Rossmann-fold superfamily protein involved in detoxification and responding to reactive carbonyl species (Ksas et al., 2024); and a phosphoethanolamine N-methyltransferase (PEAM1) involved in phosphatidylcholine biosynthesis (OG0000461) (Liu et al., 2019). The remaining four genes have diverse functions relating to DNA repair (OG0002653), telomere protection (OG0002585), signalling (OG0002789) and a mildew locus O (MLO)-like protein 1 OG0000548) linked to abiotic stress tolerance (Acevedo-Garcia et al., 2014).

**Figure 3.**
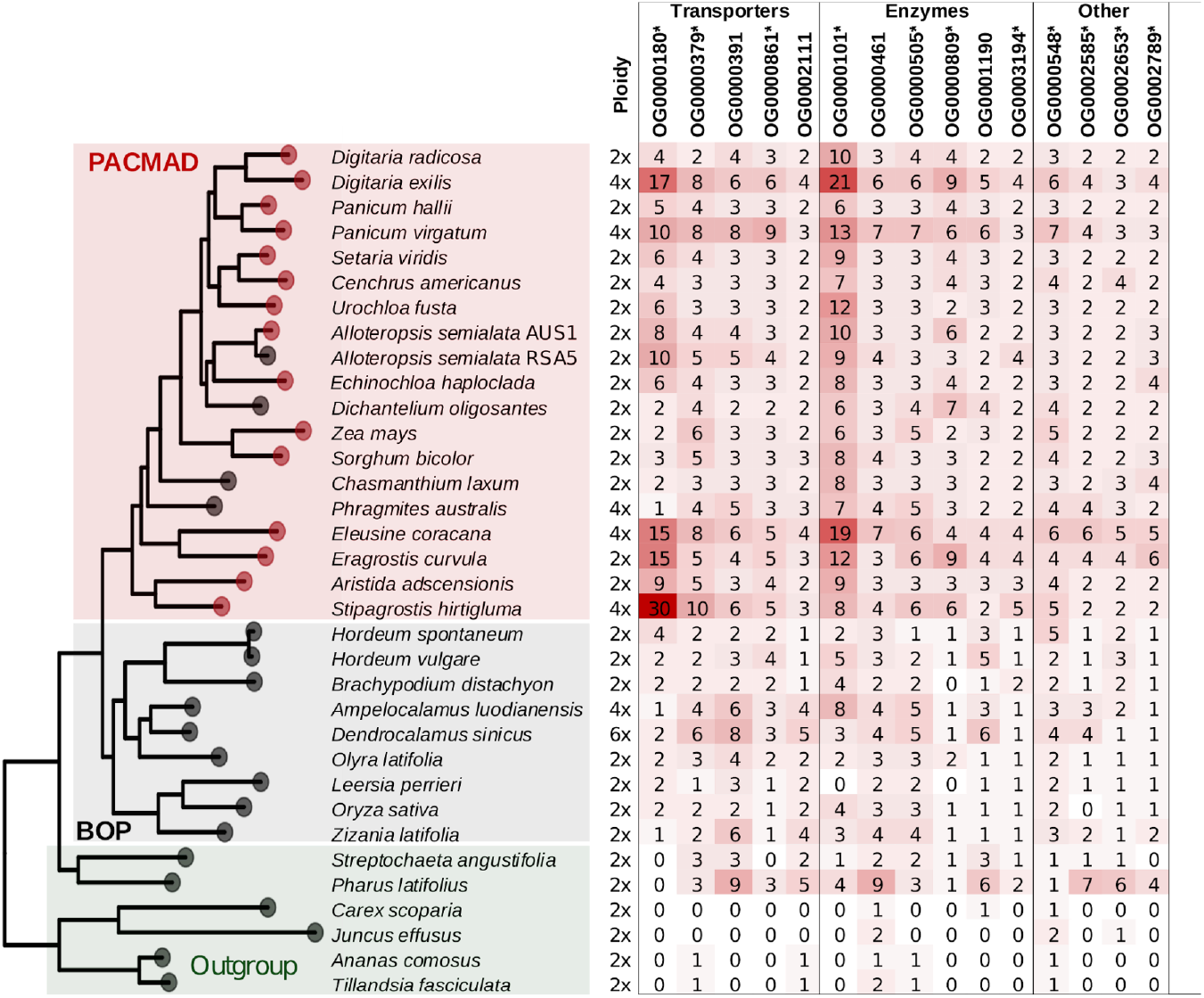
Gene copy number for each of the orthogroups that contain one of the 15 validated duplication events that occurred at the base of the PACMAD lineage and subsequently retained in all C_4_ species (these orthogroups may contain additional duplications and/or losses). Ploidy is also indicated, and the asterisks denote orthogroups which have a significantly higher gene copy number in the PACMAD than BOP clade (adjusted P-value < 0.05).

Given their reduced taxonomic distribution it is not surprising that many of the core *de novo* PACMAD genes are uncharacterised proteins (31%) or lack meaningful descriptions (Table S8). The annotated genes encode a number of transcription factors including: NAC domain-containing proteins (OG0004173), homeodomain proteins (OG0005708), ethylene-responsive transcription factors (OG0009413), response regulators (OG0016229), LNK proteins (OG0018191), FRIGIDA-like proteins (OG0018794), a WRKY domain-containing protein (OG0020072), and an orphans transcription factor (OG0019167). There are also a few core *de novo* transporters that move lipids (OG0019414) and small solutes across cell membranes in response to chemiosmotic gradients (OG0020033). There were only six core BOP orthogroup losses retained in all 15 C_4_ species, five with annotations (Table S8). These genes encoded a protein-serine/threonine phosphatase (OG0014342), lipid transporter (OG0019164), microtubule regulatory protein (OGOO18324), cupredoxin superfamily protein that catalyses inter-molecular electron transfer reactions (OGOO18546), and a membrane attack complex/perforin (MACPF) pathogen resistance protein (OG0017688).

### Relatively few known C_4_ genes identified as precursors

Among the key C_4_ metabolic enzymes, only a gene encoding β-carbonic anhydrase (OG0001853) was a candidate C_4_ precursor, β-carbonic anhydrase is the entry step of the C_4_ cycle where it is responsible for the initial fixation of CO_2_ into bicarbonate. The duplication corresponds to the specific gene lineage co-opted for C_4_ photosynthesis in grasses (β-carbonic anhydrase 2P3; Moreno-Villena et al., 2018), and although it was duplicated at the base of the PACMAD clade as previously hypothesised (Besnard et al., 2018), both duplicates were not retained in all C_4_ species (Fig. 4). Another potential direct link to the evolution of C_4_ photosynthesis could involve enzymes that are part of the photorespiration cycle. A gene encoding (DL)-glycerol-3-phosphatase (OG0003194) was duplicated at the base of PACMAD, with both copies retained in all C_4_ species used in this study. This enzyme is considered a putative 2-phosphoglycolate phosphatase (PGLP) as it shows homology to prokaryotic PGLPs and therefore may play a role in catalysing the first step of photorespiration, although this has not been functionally verified (Schwarte and Bauwe, 2007).

**Figure 4.**
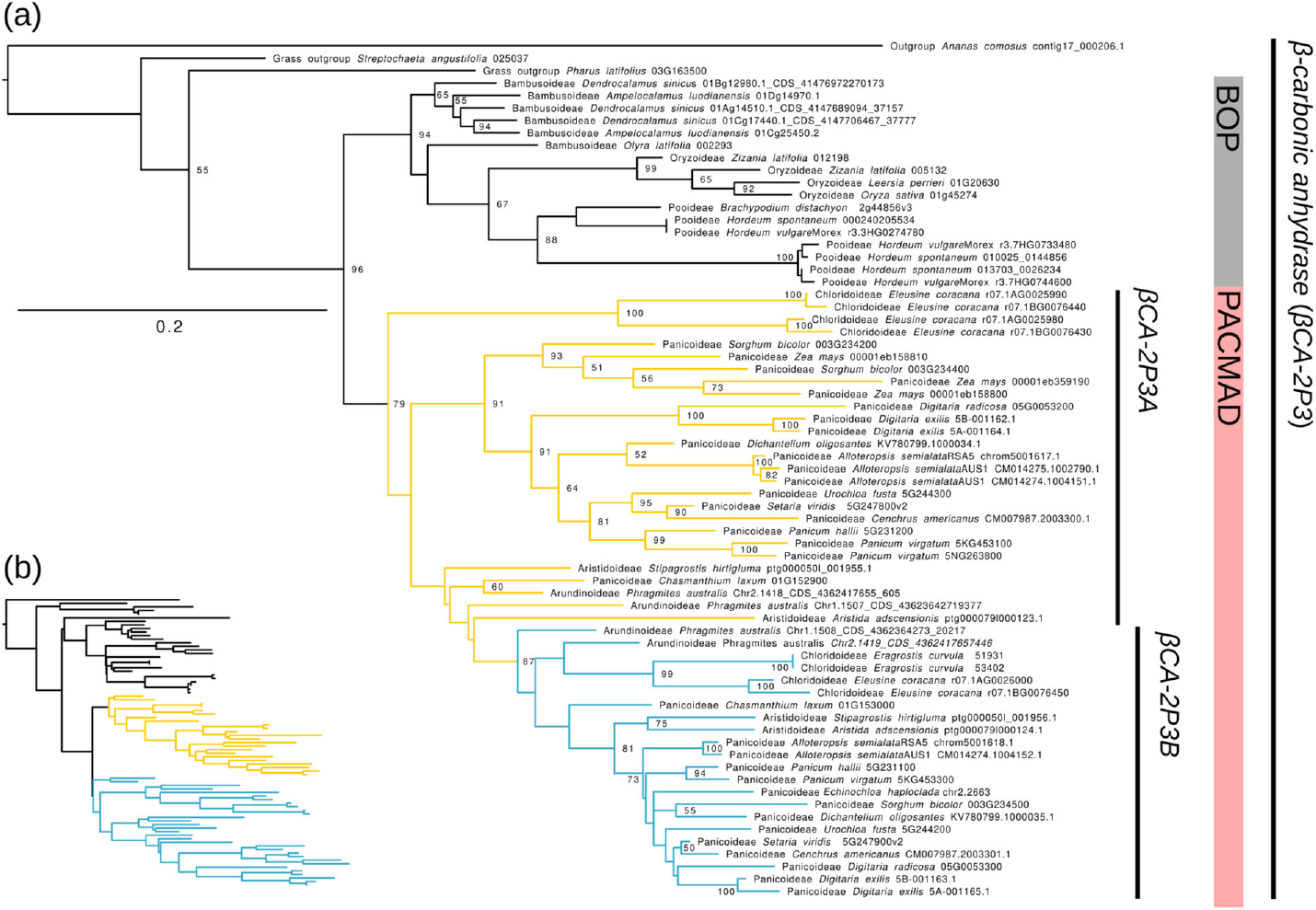
Phylogenetic trees showing the PACMAD duplication of //-carbonic anhydrase 2P3 *(βCA-2P3)*. (a) A maximum likelihood phylogeny with bootstrap support values ≥50% shown; (b) OrthoFinder-resolved gene tree (OG0001853).

### Differential gene expression in mesophyll and bundle sheath cells

In maize, 10.6% of the 32,540 gene models were significantly differentially expressed between M and BS cells (Li et al., 2010), while in sorghum it was 4.9% of the 34,211 genes (Döring et al., 2016). There was no significant difference in the proportion of genes differentially expressed between M and BS in our list of PACMAD-specific duplicates compared to the genome-wide levels in sorghum (1,035 duplicates, 54 [5.2%] differentially expressed, p-value = 0.789) (Table S9), and a significant decrease in our maize candidates (917 duplicates, 66 [7.2%] differentially expressed, p-value = 0.003) (Table S10). There was also no significant difference in the proportion of genes differentially expressed between M and BS when only considering the core PACMAD duplicates retained in all C_4_ species compared to the genome-wide proportion in both sorghum (45 duplicates, 4 [8.9%] differentially expressed, p-value = 0.261) and maize (40 duplicates, 5 [12.5%] differentially expressed, p-value = 0.890).

There were 14 PACMAD-specific duplicates that contained at least one ortholog that was significantly differentially expressed between M and BS cells in both maize and sorghum, 4 of which were core candidates: PEAM1 (OG0000461), MLO1 (OG0000548), PGML (OG0001190) and SALR (OGOOOOIOI) (Table 3). Over half (n = 8) of the differentially-expressed gene duplicates shared between maize and sorghum were not differentially expressed in rice (Table 3 & Table Sil). The remaining six were differentially expressed in all three species, two of which were more highly expressed in the same tissue (including β-carbonic anhydrase [OG0001853]), and two with opposite tissue-specific expression patterns between rice and maize + sorghum (including PEAM1 [OG0000461] and PGML [OG0001190]).

**Table 3.**
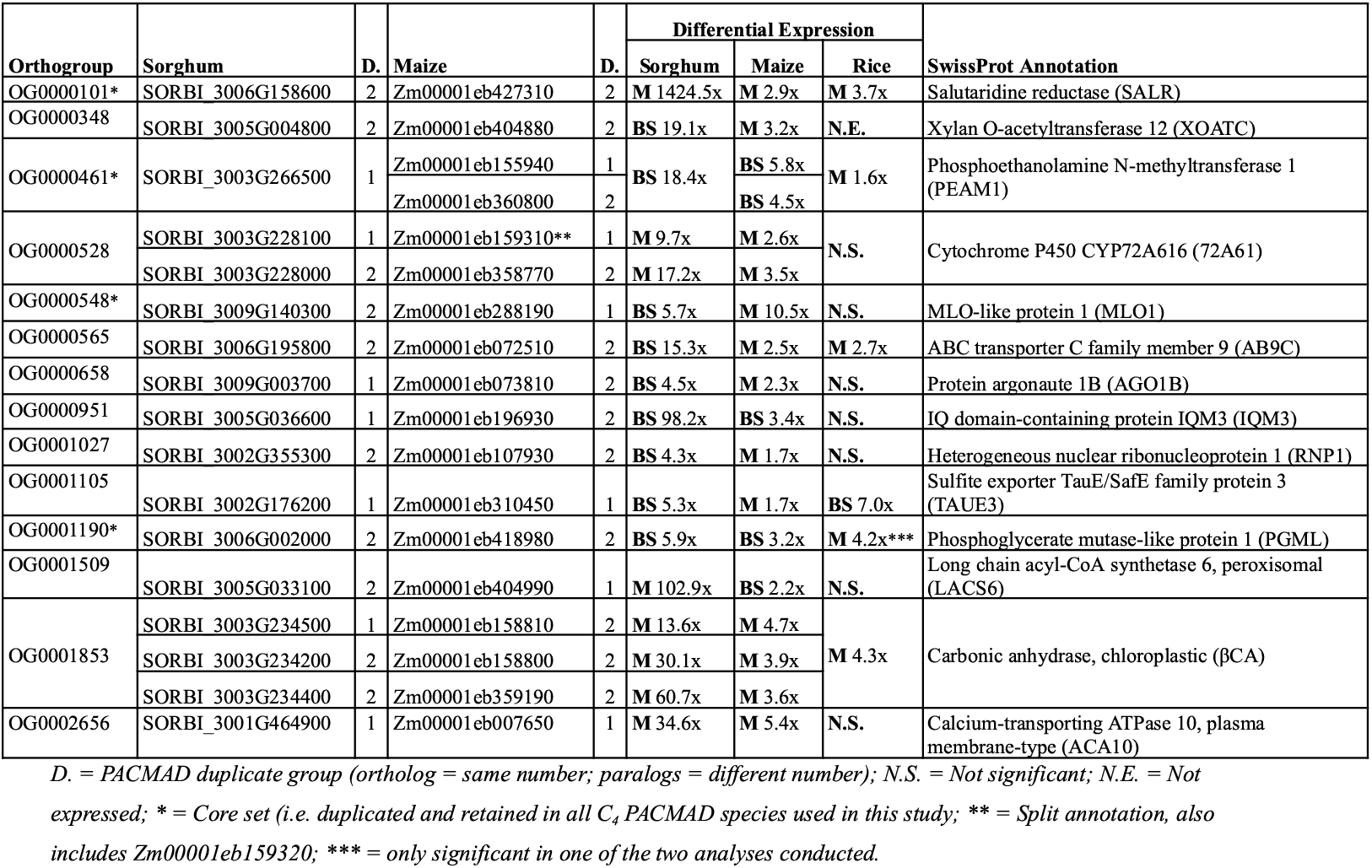
PACMAD-specific duplicates that contain at least one maize and sorghum ortholog that is significantly differentially expressed between mesophyll (M) and bundle sheath (BS) cells. The corresponding expression pattern for rice ortholog is also shown.

## Discussion

C_4_ photosynthesis is a remarkable example of convergent evolution, yet the numerous C_4_ origins are clustered in a relatively small number of major families and infrafamily groups (Christin et al., 2015). In grasses, 99% of species are found in two roughly equally size clades (BOP = 5,941 species; PACMAD = 5,815 species; Soreng et al., 2022), and the >20 origins of C_4_ in this family are restricted to the PACMAD clade (Grass Phylogeny Working Group II, 2012). Here, we assembled two genomes for the Aristidoideae {*Aristida adscensionis* and *Stipagrostis hirtigluma*}, a subfamily of grasses that includes over 350 species and is the sister subfamily to all other PACMAD grasses. We then used comparative genomics to identify genes that were duplicated or originated at the base of PACMAD or lost in the BOP clade, since these potentially represent genetic precursors underlying the repeated evolution of C_4_ physiology.

### No increase in gene duplication or subfimctionalization at the base of PACMAD

A significant increase in the rate of gene duplication is one explanation as to why genetic precursors arose specifically in the PACMAD lineage. However, there is no evidence for a whole genome duplication event at the base of this clade (Zhang et al., 2024), nor do we detect a significantly increased duplication rate along this branch. Furthermore, genes duplicated at the base of PACMAD show no increase in subfunctionalization, with a similar proportion of differential expression between C_4_ photosynthetic tissues (mesophyll and bundle sheath) to genome-wide levels. However, duplication and subfunctionalization may still play a role in the evolution of C_4_ photosynthesis in grasses, with specific examples identified in this (Table 3) and previous (Emms et al., 2016) studies. The PACMAD-specific gene duplications supplement existing variation to create a pool of paralogues that can later be independently recruited. This means the initial duplication events that are foundational for the later emergence of the trait were not directly driven by selection for a C_4_ function. A similar scenario is observed in *Solanum* species where paralog diversification generates the genetic substrate and contingency for convergent crop trait evolution (Benoit et al., 2025).

### Duplication of β-carbonic anhydrase and carbon metabolism enzymes

The evolution of C_4_ photosynthesis involves co-opting the functions of numerous enzymes to increase the efficiency and rate of carbon fixation in high temperature and light environments. Duplication and neofunctionalization is not thought to give rise directly to C_4_ enzymes (Gowik and Westhoff 2011; Williams et al. 2012; van den Bergh et al. 2014). However, older duplications in a gene family can result in paralogs that diverge in expression levels and patterning through drift or selection. The likelihood of recruiting the C_4_ specific copy then depends on the ancestral expression patterns and catalytic properties that are closest to the C_4_ optima (Wang et al., 2009; Hibberd and Covshoff, 2010; Aubry et al., 2011; Christin et al., 2013a, 2015; Emms et al., 2016; Moreno-Villena et al., 2018; Dunning et al., 2019). In the last common ancestor of the PACMAD grasses, only a single gene encoding a core C_4_ enzyme was duplicated: β-carbonic anhydrase (βCA).

βCA accounts for up to 1-2% of soluble leaf protein in higher plants (Tiwari et al., 2005), and it catalyses the first step of the C_4_ cycle where CO_2_ is initially fixed into bicarbonate in the mesophyll cells. While βCA was originally thought to play a role in the ancestral C_3_ photosynthetic pathway, CRISPR mutants in tobacco have shown that βCA is not required for C_3_ photosynthesis but instead plays a crucial role in leaf development (Hines et al., 2021). βCA is encoded by three main paralogs in grasses *(βCA-lPl, βCA-2P2 & βCA-2P3)*, with only *βCA-2P3* recruited for C_4_ photosynthesis (Moreno-Villena et al., 2018). The PACMAD-specific duplicate is of the C_4_ *βCA-2P3* copy, with the availability of reference genomes from Aristidoideae confirming that this happened in the last common ancestor of the PACMAD clade and not during the early diversification of this clade (Besnard et al., 2018). The duplication of this gene may have later facilitated independent neofunctionalization of a C_4_ copy without the need to alter its ancestral mesophyll specificity (Table 3). However, not all C_4_ species within PACMAD have retained both duplicates. One of the *βCA-2P3* daughter lineages has been lost independently in *Echinochloa haploclada* and maize (Fig. 4), although in the latter case, the remaining lineage has undergone multiple independent duplications, potentially again highlighting the role of genetic contingency in convergent trait evolution.

The duplication of other carbon metabolism enzymes may indirectly facilitate the emergence of C_4_ photosynthesis. The evolution of C_4_ photosynthesis is thought to be preceded by the establishment of a photorespiratory CO_2_ pump that is partitioned between cell types (Bräutigam and Gowik, 2016). In *Flaveria* (Asteraceae), a gene duplication facilitated the establishment of this CO_2_ pump as multiple copies enabled subfunctionalization and bundle sheath specificity (Schulze et al., 2013). Potentially a similar evolutionary scenario could occur in the PACMAD grasses which all possess a duplicated gene encoding (DL)-glycerol-3-phosphatase (OG0003194). This enzyme is considered a putative PGLP, a key enzyme of photorespiration, although this has not been functionally verified (Schwarte and Bauwe, 2007). The C_4_ and C_3_ photosynthetic pathways are also able to share carbon flux by the interchange of phosphoenolpyruvate (PEP) and 3-phosphoglycerate using the phosphoglycerate mutase and enolase pathway (Furbank and Leegood, 1984; von Caemmerer and Furbank, 2016; Furbank and Kelly, 2021). A phosphoglycerate mutase-like (PGML) protein was duplicated at the base of the PACMAD clade, and is retained in all C_4_ species (Table S3) and one of the daughter copies has become bundle sheath-specific (Table 3). The duplication of PGML and other carbon metabolism enzymes (Table S3) may generate the genetic substrate facilitating the spatial partitioning of carbohydrate metabolism in C_4_ species, with sucrose predominantly synthesised in the mesophyll and starch in the bundle sheaths (Furbank and Kelly, 2021), although future studies are needed to confirm the functional consequences of these duplications.

### Repeated duplications of amino acid and nitrate transmembrane transporters

Intercellular transporters are responsible for shuttling key metabolites between the mesophyll and bundle sheath tissues in C_4_ photosynthetic species. There has been progress in identifying many of the associated transporters, but there are still major gaps in our understanding (Mattinson and Kelly, 2025). Genes duplicated at the base of the PACMAD lineage are enriched for transmembrane amino acid transporters, potentially highlighting their role as genetic precursors of C_4_ photosynthesis. This is particularly relevant as specific amino acids, such as malate and aspartate, are key substrates of the C_4_ pathway itself. A third of the 15 core PACMAD duplicates (i.e. retained in all C_4_ species) are transporters, two of which move amino acids: cationic amino acid transporter 6 (CAAT6 - OG0002111) and organic cation/camitine transporter 7 (OG0000861). The remaining transmembrane transporters include two plant nitrate transporters in the NRT1/ PTR family (PTR15 [OG0000391] and PTR18 [OG0000379]). In C_3_-C_4_ intermediate plants that operate a glycine shuttle there is a nitrogen imbalance caused by the release of ammonia (NH_3_) in the bundle sheath. Indeed, this potential problem has been cited as one of the reasons why intermediate species might rapidly transition to a full C_4_ cycle (Mailman et al., 2014; Bräutigam and Gowik, 2016). The duplicated nitrate and amino acid transporters may play a role in mitigating the nitrogen and amino-group imbalance during the incipient stages of C_4_ evolution.

### Enhancing defence against oxidative stress in bundle sheath cells

Oxygenic photosynthesis generates abundant reactive oxygen species (ROS). While ROS can play key signalling roles, they can also damage cells and impair photosynthesis. To maintain redox homeostasis plants have evolved several enzymatic and non-enzymatic strategies for ROS (Turkan et al., 2018). Glutathione is one of the key cellular pools of antioxidants (Mittler et al., 2004), and genes duplicated at the base of PACMAD are enriched for glutathione transferase activity (G0:0004364) and glutathione metabolic process (G0:0006749). The evolutionary trajectory to C_4_ should overall reduce photorespiration and ROS production, but the dynamics of ROS production in the different leaf compartments require a tailored solution (Turkan et al., 2018). Comparisons between different C_4_ and C_3_ species show that regulatory and redox-based systems appear to be quite variable and are likely to be specific to the decarboxylation enzyme used by the plant for C_4_ photosynthesis (Bräutigam et al., 2014; Turkan et al., 2018).

Out of the 15 core PACMAD duplicates, three have functions directly relating to limiting oxidative damage: a peroxidase (OG0000809), SALR (OGOOOOIOI) and PEAM1 (OG0000461). The duplication of phosphoethanolamine N-methyltransferase 1 (PEAM1) is correlated with an expression shift from being mesophyll (C_3_ rice) to bundle sheath (C_4_ maize and sorghum) specific (Table 3). The same gene was also identified in a genome-wide association study (GWAS) in *Alloteropsis semialata* for the inner bundle sheath fraction (Alenazi et al., 2024), which is the proportion of the leaf responsible for refixing carbon through the C_4_ cycle (Alenazi et al., 2023). PEAM1 catalyses the synthesis of phosphatidylcholine, a major membrane phospholipid (Chen et al., 2018), and orthologs in *Arabidopsis* are involved in regulating salt tolerance (He et al., 2022) and root cell differentiation (Zou et al., 2019) by balancing ROS production and abscisic acid or auxin, respectively. SALR (OGOOOOIOI) is mesophyll-specific (Table 3) and orthologous to an Arabidopsis NAD(P)-binding Rossmann-fold superfamily protein involved in detoxification and responding to reactive carbonyl species (Ksas et al., 2024). The repeated duplication of genes associated with oxidative stress at the base of PACMAD created redundancy in the genetic basis of oxidative stress management. This contingency enabled the selective recruitment and subfunctionalization of loci to streamline oxidative stress management for a C_4_ context.

### Multiple PACMAD-specific transcription regulators

Genes co-opted for C_4_ photosynthesis become highly expressed in leaf tissue, but it still remains unclear how many of these are regulated, particularly for trans-acting factors (Hibberd and Covshoff, 2010; Schlüter and Weber, 2020; Lyu et al., 2025). A recent study on the evolution of C_4_ photosynthesis in *Flaveria* showed that the emergence of this trait was correlated with the expansion of ethylene response factor (ERF) transcription factors (Lyu et al., 2025). In PACMAD we identify a number of *de novo* transcription factor orthogroups, including a group of ERFs retained in all C_4_ lineages (OG0009413). One possible explanation why C_4_ photosynthesis is restricted to the PACMAD clade is the *de novo* origin of transcription factors that could be readily recruited for this process.

## Conclusion

The assembly of the first genomes for species belonging to the subfamily Aristidoideae permits the inference of genetic changes in the last common ancestor of the PACMAD clade that enabled the repeated evolution of C_4_ photosynthesis. Gene duplication, *de novo* formation and loss has created the prerequisite standing variation in gene content that could later be co-opted for C_4_ photosynthesis in the PACMAD clade. This includes an additional copy of a core C_4_ enzyme (β-carbonic anhydrase), as well as genes with C_4_ related functions including carbon metabolism, amino acid and nitrate transport, oxidative stress management and transcription regulation. This gene turnover created the necessary genetic contingency, facilitating the independent recruitment and refinement of C_4_ genes in the independent origins. This supports the idea that the repeated evolution of C_4_ photosynthesis in PACMAD grasses was not driven by a single genetic event, but rather by the opportunistic utilisation of pre-existing genetic diversity.

## Data and code availability statement

The Aristidoideae raw data, genomes and annotation files are available in NCBI (XXX). Tree files and Helixer re-annotations of published genomes are available on dryad (XXX). The genome assembly and annotation code are available in GitHub (https://github.com/Sheffield-Plant-Evolutionary-Genomics).

## Author contributions

LP, PAC and LTD conceptualised and designed the study. LP generated the genome assemblies. AA and LP performed the duplication and gene loss analyses. SM and IJL obtained genome size estimates. All authors helped interpret the data. LP, AA and LTD wrote the manuscript with the help of all authors.

## Conflict of interest statement

The authors declare no conflicts of interest.

## Acknowledgements

We thank the staff of the Arthur Willis Environment Centre and all members of the Dunning and Osborne labs for their help with plant growth and maintenance. We thank Lucy Knowles for her support in DNA quality assessment. Sequencing data generation was carried out by the Centre for Genomic Research, which is based at the University of Liverpool. This work was funded by the Natural Environment Research Council (grant no.: NE/V000012/1). PAC was funded by a Royal Society University Research Fellowship (grant no.: URF\R\180022). LTD is funded by a NERC fellowship (grant no.: NE/T011025/1). The authors also extend their appreciation to the Deanship of Scientific Research at Northern Border University, Arar, KSA for funding this research work through the project number NBU-FPEJ-2025-630-01.

## Notes

### Competing Interest Statement

The authors have declared no competing interest.

## References

Acevedo-Garcia J, Kusch S, Panstruga R. 2014. Magical mystery tour. MLO proteins in plant immunity and beyond. New Phytologist 204: 273–281.

Alenazi AS, Bianconi ME, Middlemiss E, Milenkovic V, Curran EV, Sotelo G et al. 2023. Leaf anatomy explains the strength of C_4_ activity within the grass species Alloteropsis semialata. Plant, Cell & Environment 46: 2310–2322.

Alenazi AS, Pereira L, Christin P-A, Osborne CP, Dunning LT. 2024. Identifying genomic regions associated with C_4_ photosynthetic activity and leaf anatomy in Alloteropsis semialata. New Phytologist 243: 1698–1710.

Ashbumer M, Ball CA, Blake JA, Botstein D, Butler H, Cherry JM et al. 2000. Gene Ontology: tool for the unification of biology. Nature Genetics 25: 25–29.

Atkinson RRL, Mockford EJ, Bennett C, Christin P-A, Spriggs EL, Freckleton RP et al. 2016. C_4_ photosynthesis boosts growth by altering physiology, allocation and size. Nature Plants 2: 16038.

Aubry S, Brown NJ, Hibberd JM. 2011. The role of proteins in C_3_ plants prior to their recruitment into the C_4_pathway. Journal of Experimental Botany 62: 3049–3059.

Benoit M, Jenike KM, Satterlee JW, Ramakrishnan S, Gentile I, Hendelman A et al. 2025. Solanum pan-genetics reveals paralogues as contingencies in crop engineering. Nature 640: 135–145.

Berardini TZ, Reiser L, Li D, Mezheritsky Y, Muller R, Strait E, Huala E. 2015. The Arabidopsis information resource: making and mining the “gold standard” annotated reference plant genome. Genesis 53: 474–485.

Besnard G, Bianconi ME, Hackel J, Manzi S, Vorontsova MS, Christin PA. 2018. Herbarium genomics retraces the origins of C_4_-specific carbonic anhydrase in Andropogoneae (Poaceae). Botany Letters 165: 419–33.

Bianconi ME, Dunning LT, Moreno-Villena JJ, Osborne CP, Christin PA. 2018. Gene duplication and dosage effects during the early emergence of C_4_ photosynthesis in the grass genus Alloteropsis. Journal of Experimental Botany 69: 1967–1980.

Bräutigam A, Gowik U. 2016. Photorespiration connects C_3_ and C_4_ photosynthesis. Journal of Experimental Botany 6T. 2953-2962.

Bräutigam A, Schliesky S, Kŭlahoglu C, Osborne CP, Weber AP. 2024. Towards an integrative model of C_4_ photosynthetic subtypes: insights from comparative transcriptome analysis of NAD-ME, NADP-ME, and PEP-CK C_4_ species. Journal of Experimental Botany 65: 3579–93.

Bussotti G, Raineri E, Erb I, Zytnicki M, Wilm A, Beaudoing E, Bucher P, Notredame C. 2011. BlastR—fast and accurate database searches for non-coding RNAs. Nucleic Acids Research 39: 6886–6895.

Chau LM, Goodisman MAD. 2017. Gene duplication and the evolution of phenotypic diversity in insect societies. Evolution 71: 2871–2884.

Chen W, Salari H, Taylor MC, Jost R, Berkowitz O, Barrow R et al. 2018. NMT1 and NMT3 V-methyltransferase activity is critical to lipid homeostasis, morphogenesis, and reproduction. Plant Physiology 177: 1605–1628.

Chen Y, Zhang Y, Wang AY, Gao M, Chong Z. 2021. Accurate long-read de novo assembly evaluation with Inspector. Genome Biology 22: 312.

Cheng H, Concepcion GT, Feng X, Zhang H.Li H. 2021. Haplotype-resolved de novo assembly using phased assembly graphs with hifiasm. Nature Methods 18: 170–175.

Christin P-A, Arakaki M, Osborne CP, Edwards EJ. 2015. Genetic enablers underlying the clustered evolutionary origins of C_4_ photosynthesis in angiosperms. Molecular Biology and Evolution 32: 846–858.

Christin PA, Boxall SF, Gregory R, Edwards EJ, Hartwell J, Osborne CP. 2013a. Parallel recruitment of multiple genes into C_4_ photosynthesis. Genome Biology and Evolution 5: 2174–2187.

Christin PA, Osborne CP, Chatelet DS, Columbus JT, Besnard G, Hodkinson TR et al. 2013b. Anatomical enablers and the evolution of C_4_ photosynthesis in grasses. Proceedings of the National Academy of Sciences 110: 1381–1386.

Christin P-A, Spriggs E, Osborne CP, Strömberg CA, Salamin N, Edwards EJ. 2014. Molecular dating, evolutionary rates, and the age of the grasses. Systematic Biology 63: 153–165.

Clark J, Hidalgo O, Pellicer J, Liu H, Marquardt J, Robert Y, Christenhusz M et al. 2016. Genome evolution of ferns: evidence for relative stasis of genome size across the fern phylogeny. New Phytologist 210: 1072–1082.

Conant GC, Wolfe KH. 2008. Turning a hobby into a job: how duplicated genes find new functions. Nature Reviews Genetics 9: 938–950.

Darwin, C. 1859. On the Origin of Species by Means of Natural Selection. John Murray, London.

Doležel J, Greilhuber J, Suda J. 2007. Estimation of nuclear DNA content in plants using flow cytometry. Nature Protocols 2: 2233–2244.

Döring F, Streubel M, Bräutigam A, Gowik U. 2016. Most photorespiratory genes are preferentially expressed in the bundle sheath cells of the C_4_ grass Sorghum bicolor. Journal of Experimental Botany 67: 3053–3064.

Drouin G, Godin JR, Page B. 2011. The genetics of vitamin C loss in vertebrates. Current Genomics 12: 371–378.

Dunning LT, Lundgren MR, Moreno-Villena JJ, Namaganda M, Edwards EJ, Nosil P et al. 2017. Introgression and repeated co-option facilitated the recurrent emergence of C_4_ photosynthesis among close relatives. Evolution 71: 1541–55.

Dunning LT, Moreno-Villena JJ, Lundgren MR, Dionora J, Salazar P, Adams C, Nyirenda F et al. 2019. Key changes in gene expression identified for different stages of C_4_ evolution in Alloteropsis semialata. Journal of Experimental Botany 70: 3255–3268.

Edgar RC. 2994. MUSCLE: multiple sequence alignment with high accuracy and high throughput. Nucleic Acids Research. 32: 1792–7.

Ehleringer JR. 1978. Implications of quantum yield differences on the distributions of C_3_ and C_4_ grasses. Oecologia 31: 255–267.

Ehleringer J, Björkman O. 1977. Quantum yields for CO_2_ uptake in C_3_ and C_4_ plants: dependence on temperature, CO_2_, and O_2_ concentration. Plant Physiology 59: 86–90.

Ehleringer JR, Ceriing TE, Helliker BR. 1997. C_4_ photosynthesis, atmospheric CO_2_, and climate. Oecologia 112: 285–299.

Ehleringer JR, Sage RF, Flanagan LB, Pearcy RW. 1991. Climate change and the evolution of C_4_ photosynthesis. Trends in Ecology & Evolution 6: 95–99.

Emms DM, Covshoff S, Hibberd JM, Kelly S. 2016. Independent and parallel evolution of new genes by gene duplication in two origins of C_4_ photosynthesis provides new insight into the mechanism of phloem loading in C_4_ species. Molecular Biology and Evolution 33: 1796–1806.

Emms DM, Kelly S. 2017. STRIDE: species tree root inference from gene duplication events. Molecular Biology and Evolution 34: 3267–3278.

Emms DM, Kelly S. 2018 STAG: Species Tree Inference from All Genes. BioRxiv 267914; 10.1101/267914

Emms DM, Kelly S. 2019. OrthoFinder: phylogenetic orthology inference for comparative genomics. Genome Biology 20: 1–14.

Femández-Pérez F, Pomar F, Pedreño MA, Novo-Uzal E. 2015. The suppression of AtPrx52 affects fibers but not xylem lignification in Arabidopsis by altering the proportion of syringyl units. Physiologia Plantarum 154: 395–406.

Furbank RT, Kelly S. 2021. Finding the C_4_ sweet spot: cellular compartmentation of carbohydrate metabolism in C_4_photosynthesis. Journal of Experimental Botany 72: 6018–6026.

Furbank RT, Leegood RC. 1984. Carbon metabolism and gas exchange in leaves of Zea mays L. Interaction between the C_3_ and C_4_ pathways during photosynthetic induction. Planta 162: 457–462.

Gowik U, Westhoff P. 2011. The path from C_3_ to C_4_ photosynthesis. Plant physiology 155: 56–63.

Grass Phylogeny Working Group II. 2012. New grass phylogeny resolves deep evolutionary relationships and discovers C_4_ origins. New Phytologist 193: 304–312.

Guindon S, Dufayard JF, Lefort V, Anisimova M, Hordijk W, Gascuel O. 2010. New algorithms and methods to estimate maximum-likelihood phytogenies: assessing the performance of PhyML 3.0. Systematic Biology 59: 307–321.

Hatch MD. 1987. C_4_ photosynthesis: a unique elend of modified biochemistry, anatomy and ultrastructure. Biochìmica et Biophysica Acta (BBA) - Reviews on Bioenergetics 895: 81–106.

He QY, Jin JF, Lou HQ, Dang FF, Xu JM et al. 2022. Abscisic acid-dependent PMT1 expression regulates salt tolerance by alleviating abscisic acid-mediated reactive oxygen species production in Arabidopsis. Journal of Integrative Plant Biology 64: 1803–1820.

Helsen J, Voordeckers K, Vanderwaeren L, Santermans T, Tsontaki M, Verstrepen KJ, Jelier R. 2020. Gene toss predictably drives evolutionary adaptation. Molecular Biology and Evolution 37: 2989–3002.

Heyduk K, Moreno-Villena JJ, Gilman IS, Christin PA, Edwards EJ. 2019. The genetics of convergent evolution: insights from plant photosynthesis. Nature Reviews Genetics 20: 485–493.

Hibberd JM, Covshoff S. 2010. The regulation of gene expression required for C_4_ photosynthesis. Annual Review of Plant Biology 61: 181–207.

Hines KM, Chaudhari V, Edgeworth KN, Owens TG, Hanson MR. 2021, Absence of carbonic anhydrase in chloroplasts affects C_3_ plant development but not photosynthesis. Proceedings of the National Academy of Sciences 118:e2107425118.

Hoang CF, Liu WY, Lu MYJ, Chen YH, Ku MS, Li WH. 2021. Whole-genome duplication facilitated the evolution of C_4_ photosynthesis in Gynandropsis gynandra. Molecular Biology and Evolution 38: 4715–4731.

Holst F, Bolger A, Gŭnther C, Maβ J, Triesch S, Kindel F et al. 2023. Helixer-de novo prediction of primary eukaryotic gene models combining deep learning and a hidden Markov model. BioRxiv 2023-02.

Howe KL, Contreras-Moreira B, De Silva N, Maslen G, Akanni W, Allen J et al. 2020. Ensembl Genomes 2020—enabling non-vertebrate genomic research. Nucleic Acids Research 48: D689–D695.

Hua L, Stevenson SR, Reyna-Llorens I, Xiong H, Kopriva S, Hibberd JM. 2021. The bundle sheath of rice is conditioned to play an active role in water transport as well as sulfur assimilation and jasmonic acid synthesis. The Plant Journal 107: 268–286.

Hunt M, Silva ND, Otto TD, Parkhill J, Keane JA, Harris SR. 2015. Circlator: automated circularization of genome assemblies using long sequencing reads. Genome Biology 16: 294.

Huu CN, Keller B, Conti E, Kappel C, Lenhard M. 2020. Supergene evolution via stepwise duplications and neofunctionalization of a floral-organ identity gene. Proceedings of the National Academy of Sciences MT. 23148–23157.

Jiang M, Zhan Z, Li H, Dong X, Cheng F, Piao Z. 2020. Brassica rapa orphan genes largely affect soluble sugar metabolism. Horticulture Research 7.

Katoh K, Standley DM. 2013. MAFFT multiple sequence alignment software version 7: improvements in performance and usability. Molecular Biology and Evolution 30: 772–780.

Ksas B, Chiarenza S, Dubourg N, Menard V, Gilbin R, Havaux M. 2024. Plant acclimation to ionising radiation requires activation of a detoxification pathway against carbonyl-containing lipid oxidation products. Plant, Cell & Environment 47: 3882–3898.

Lefort V, Longueville JE, Gascuel O. 2017. SMS: smart model selection in PhyML. Molecular Biology and Evolution, 34: 2422–2424.

Lenski RE, Ofria C, Pennock RT, Adami C. 2003. The evolutionary origin of complex features. Nature 423: 139–144.

Lewis MW, Hake S. 2016. Keep on growing: building and patterning leaves in the grasses. Current Opinion in Plant Biology 29: 80–86.

Li H. 2018. Minimap2: pairwise alignment for nucleotide sequences. Bioinformatics 34: 3094–3100.

Li L, Foster CM, Gan Q, Nettleton D, James MG, Myers AM, Wurtele ES. 2009. Identification of the novel protein QQS as a component of the starch metabolic network in Årabidopsis leaves. The Plant Journal 58: 485–498.

Li P, Ponnala L, Gandotra N, Wang L, Si Y, Tausta SL et al. 2010. The developmental dynamics of the maize leaf transcriptome. Nature Genetics 42: 1060.

Liu YC, Lin YC, Kanehara K, Nakamura Y. 2019. A methyltransferase trio essential for phosphatidylcholine biosynthesis and growth. Plant Physiology 179: 433–445.

Loureiro J, Rodriguez E, Doležel J, Santos C. 2007. Two new nuclear isolation buffers for plant DNA flow cytometry: a test with 37 species. Annals of Botany 100: 875–888.

Lyu MJA, D. H, Yao H, Zhang Z, Chen G, Huang Y et al. 2025. A dominant role of transcriptional regulation during the evolution of C_4_ photosynthesis in Flaveria species. Nature Communications 16: 1643.

Magadum S, Banerjee U, Murugan P, Gangapur D, Ravikesavan R. 2013. Gene duplication as a major force in evolution. Journal of Genetics 92: 155–161.

Mallmann J, Heckmann D, Bräutigam A, Lercher MJ, Weber AP, Westhoff P, Gowik U. 2014. The role of photorespiration during the evolution of C_4_ photosynthesis in the genus Flaveria. Elife 3: e02478.

Mattinson O, Kelly S. 2025. The metabolite transporters of C_4_ photosynthesis. The Plant Cell koaf019.

Meléndez-Hevia E, Waddell TG, Cascante M. 1996 The puzzle of the Krebs citric acid cycle: Assembling the pieces of chemically feasible reactions, and opportunism in the design of metabolic pathways during evolution. Journal of Molecular Evolution 43: 293–303.

Mi H, Muruganujan A, Thomas PD. 2012. PANTHER in 2013: modelling the evolution of gene function, and other gene attributes, in the context of phylogenetic trees. Nucleic Acids Research 41: D377–D386.

Mittler R, Vanderauwera S, Gollery M, Van Breusegem F. 2004. Reactive oxygen gene network of plants. Trends in Plant Science 9: 490–498.

Monson RK. 1999. The origins of C_4_ genes and evolutionary pattern in the C_4_ metabolic phenotype. C_4_ plant biology, Academic Press

Monson RK. 2003. Gene Duplication, Neofunctionalization, and the Evolution of C_4_ Photosynthesis. International Journal of Plant Sciences 164: S43–S54.

Moore, R. C., and Purugganan, M. D. (2005). The evolutionary dynamics of plant duplicate genes. Current Opinion in Plant Biology 8: 122–128.

Moreno-Villena JJ, Dunning LT, Osborne CP, Christin PA. 2018. Highly expressed genes are preferentially co-opted for C_4_ photosynthesis. Molecular Biology and Evolution 35: 94–106.

Obermayer R, Leitch IJ, Hanson L, Bennett MD. 2002. Nuclear DNA C-values in 30 species double the familial representation in pteridophytes. Annals of Botany 90: 209–217.

Ou S, Su W, Liao Y, Chougule K, Agda JRA, Hellinga AJ et al. (2019). Benchmarking transposable element annotation methods for creation of a streamlined, comprehensive pipeline. Genome Biology 20: 275.

Ourisson G, Nakatani Y. 1994 The terpenoid theory of the origin of cellular life: the evolution of terpenoids to cholesterol. Chemistry & Biology 1: 11–23.

PanchyN, Lehti-ShiuM, Shiu SH. 2016. Evolution of gene duplication in plants. Plant Physiology 171: 2294–2316.

Paparelli E, Gonzali S, Parlanti S, Novi G, Giorgi FM, Licausi F et al. 2012. Misexpression of a chloroplast aspartyl protease leads to severe growth defects and alters carbohydrate metabolism in Arabidopsis. Plant physiology 160: 1237–1250.

Pereira L, Bianconi ME, Osborne CP, Christin PA, Dunning LT. 2023. Alloteropsis semialata as a study system for C_4_ evolution in grasses. Annals of Botany 132: 365–382.

Planta J, Liang YY, Xin H, Chansler MT, Prather LA, Jiang N et al. 2022. Chromosome-scale genome assemblies and annotations for Poales species Carex cristatella, Carex scoparia, Juncus effusus, and Juncus inflexus. G3 12:jkac211.

Price MN, Dehal PS, Arkin AP. 2010. FastTree 2-approximately maximum-likelihood trees for large alignments. PloS One 5: e9490.

Quinlan AR, Hall IM. 2010. BEDTools: a flexible suite of utilities for comparing genomic features. Bioinformatics 26: 841–842.

Ranallo-Benavidez TR, Jaron KS. Schatz MC. 2020. GenomeScope 2.0 and Smudgeplot for reference-free profiling of polyploid genomes. Nature Communications 11: 1432.

Rice P, Longden I, Bleasby A. 2000. EMBOSS: the European molecular biology open software suite. Trends in Genetics 16: 276–277.

Sage RF, Christin PA, Edwards E J. 2011. The C_4_ plant lineages of planet Earth. Journal of Experimental Botany 61: 3155–3169.

Sage RF, Sage TL, Kocacinar F. 2012. Photorespiration and the evolution of C_4_ photosynthesis. Annual Review of Plant Biology 63: 19—47.

Schlüter U, Weber AP. 2020. Regulation and evolution of C_4_ photosynthesis. Annual Review of Plant Biology 71: 183–215.

Schulze S, Mallmann J, Burscheidt J, Koczor M, Streubel M et al. (2013) Evolution of C_4_ photosynthesis in the genus Flaveria. establishment of a photorespiratory CO_2_ pump. The Plant Cell 25: 2522–2535.

Schwarte S, Bauwe H. 2007. Identification of the photorespiratory 2-phosphoglycolate phosphatase, PGLP1, in Arabidopsis. Plant Physiology 144: 1580–1586.

Sedelnikova OV, Hughes TE, Langdale JA. 2018. Understanding the genetic basis of C_4_ Kranz anatomy with a view to engineering C_3_ crops. Annual Review of Genetics 52: 249–270.

Simão FA, Waterhouse RM, Ioannidis P, Kriventseva EV, Zdobnov EM. 2015. BUSCO: assessing genome assembly and annotation completeness with single-copy orthologs. Bioinformatics 31: 3210–3212.

Sloan DB, Alverson AJ, Chuckalovcak JP, Wu M, McCauley DE, Palmer JD, Taylor DR. 2012. Rapid evolution of enormous, multichromosomal genomes in flowering plant mitochondria with exceptionally high mutation rates. PLoSBiology 10: el001241.

Soreng RJ, Peterson PM, Zuloaga FO, Romaschenko K, Clark L et al. 2022. A worldwide phylogenetic classification of the Poaceae (Gramineae) III: An update. Journal of Systematics and Evolution 60: 476–521.

Spriggs EL, Christin P-A, Edwards EJ. 2014. C_4_ photosynthesis promoted species diversification during the Miocene grassland expansion. PloS One 9: e97722.

Stiehler F, Steinbom M, Scholz S, Dey D, Weber AP, Denton AK. 2020. Helixer: cross-species gene annotation of large eukaryotic genomes using deep learning. Bioinformatics 36: 5291–5298.

Thomas PD, Ebert D, Muruganujan A, Mushayahama T, Albou L, Mi H. 2021. PANTHER: Making genome-scale phylogenetics accessible to all. Protein Science 31: 8–22.

Tillich M, Lehwark P, Pellizzer T, Ulbricht-Jones ES, Fischer A, Bock R, Greiner S. 2017. GeSeq - versatile and accurate annotation of organelle genomes. Nucleic Acids Research 45: W6–W11.

Tiwari A, Kumar P, Singh S, and Ansari SA. 2005. Carbonic anhydrase in relation to higher plants. Photosynthetica 43: 1–11.

Turkan I, Uzilday B, Dietz KJ, Bräutigam A, Ozgur R. 2018. Reactive oxygen species and redox regulation in mesophyll and bundle sheath cells of C_4_ plants. Journal of Experimental Botany 69: 3321–3331.

Uliano-Silva M, Ferreira JGRN, Krasheninnikova K, Darwin Tree of Life Consortium etal. 2023. MitoHiFi: a python pipeline for mitochondrial genome assembly from PacBio high fidelity reads. BMC Bioinformatics 24: 288.

van den Bergh E, Kŭlahoglu C, Bräutigam A, Hibberd JM et al. 2014. Gene and genome duplications and the origin of C_4_ photosynthesis: birth of a trait in the Cleomaceae. Current Plant Biology 1: 2–9.

von Caemmerer S, Furbank RT. 2016. Strategies for improving C_4_ photosynthesis. Current Opinion in Plant Biology 31: 125–134.

Wagner GP, Amemiya C, Ruddle F. 2003. Hox cluster duplications and the opportunity for evolutionary novelties. Proceedings of the National Academy of Sciences 100: 14603–14606.

Wang X, Gowik U, Tang H, Bowers JE, Westhoff P, Paterson AH. 2009. Comparative genomic analysis of C_4_photosynthetic pathway evolution in grasses. Genome Biology 10: R68.

Williams, B.P., Aubry, S., & Hibberd, J.M. 2012. Molecular evolution of genes recruited into C_4_ photosynthesis. Trends in Plant Science 17: 213–220.

Woodhouse MR, Cannon EK, Portwood JL, Harper LC, Gardiner JM et al. 2021. A pan-genomic approach to genome databases using maize as a model system. BMC Plant Biology 21: 1–10.

Xia S, Chen J, Arsala D, Emerson JJ, Long M. 2025. Functional innovation through new genes as a general evolutionary process. Nature genetics 57: 295–309.

Zhang T, Huang W, Zhang L, Li DZ, Qi J, Ma H. 2024. Phylogenomic profiles of whole-genome duplications in Poaceae and landscape of differential duplicate retention and losses among major Poaceae lineages. Nature Communications 15: 3305.

Zou Y, Zhang X, Tan Y, Huang JB, Zheng Z, Tao LZ. 2019. Phosphoethanolamine Y-methyltransferase 1 contributes to maintenance of root apical meristem by affecting ROS and auxin-regulated cell differentiation in Årabidopsis. New Phytologist 224: 258–273.

