## Supplementary Material for "Gene turnover and contingency facilitated the repeated evolution of C_4_ photosynthesis in grasses"

**Figure S1**. K-mer spectrum analysis of the *Aristida adscensionis* PacBio HiFi data.

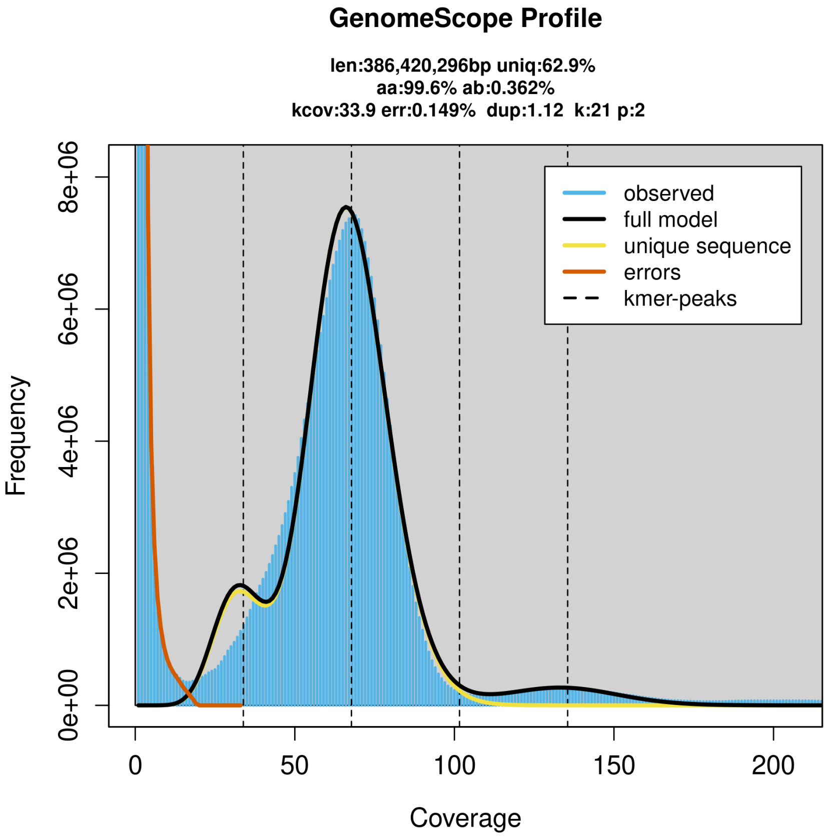

**Figure S2**. K-mer spectrum analysis of the *Stipagrostis hirtigluma* PacBio HiFi data.

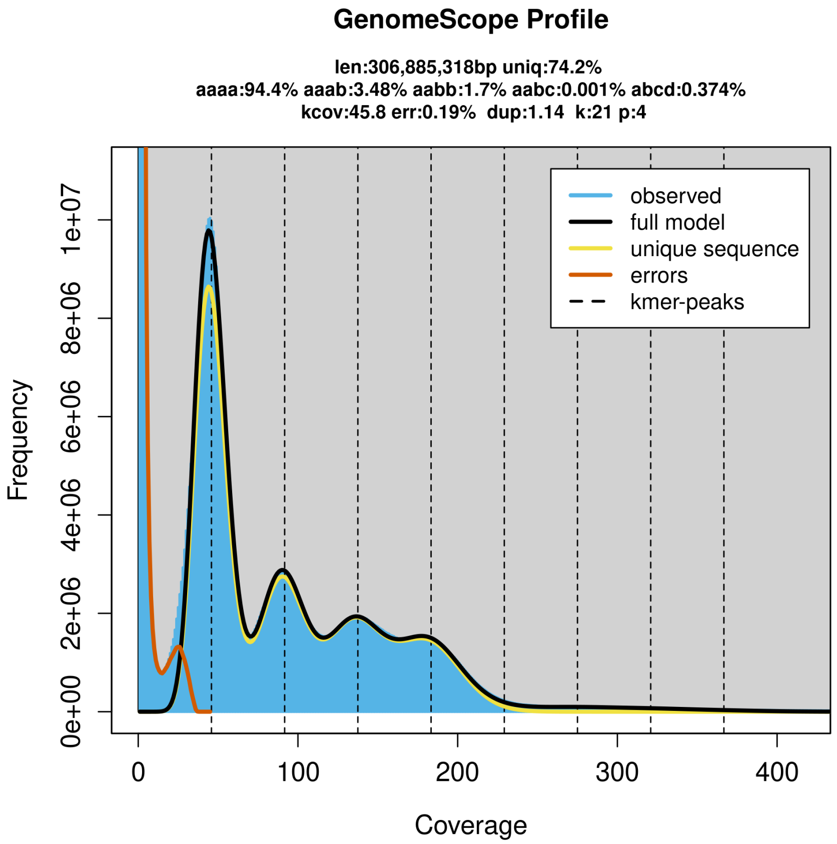

**Figure S3**. Hierarchical representation for the GO terms significantly overrepresented (in yellow boxes) in the duplicated candidate genes.

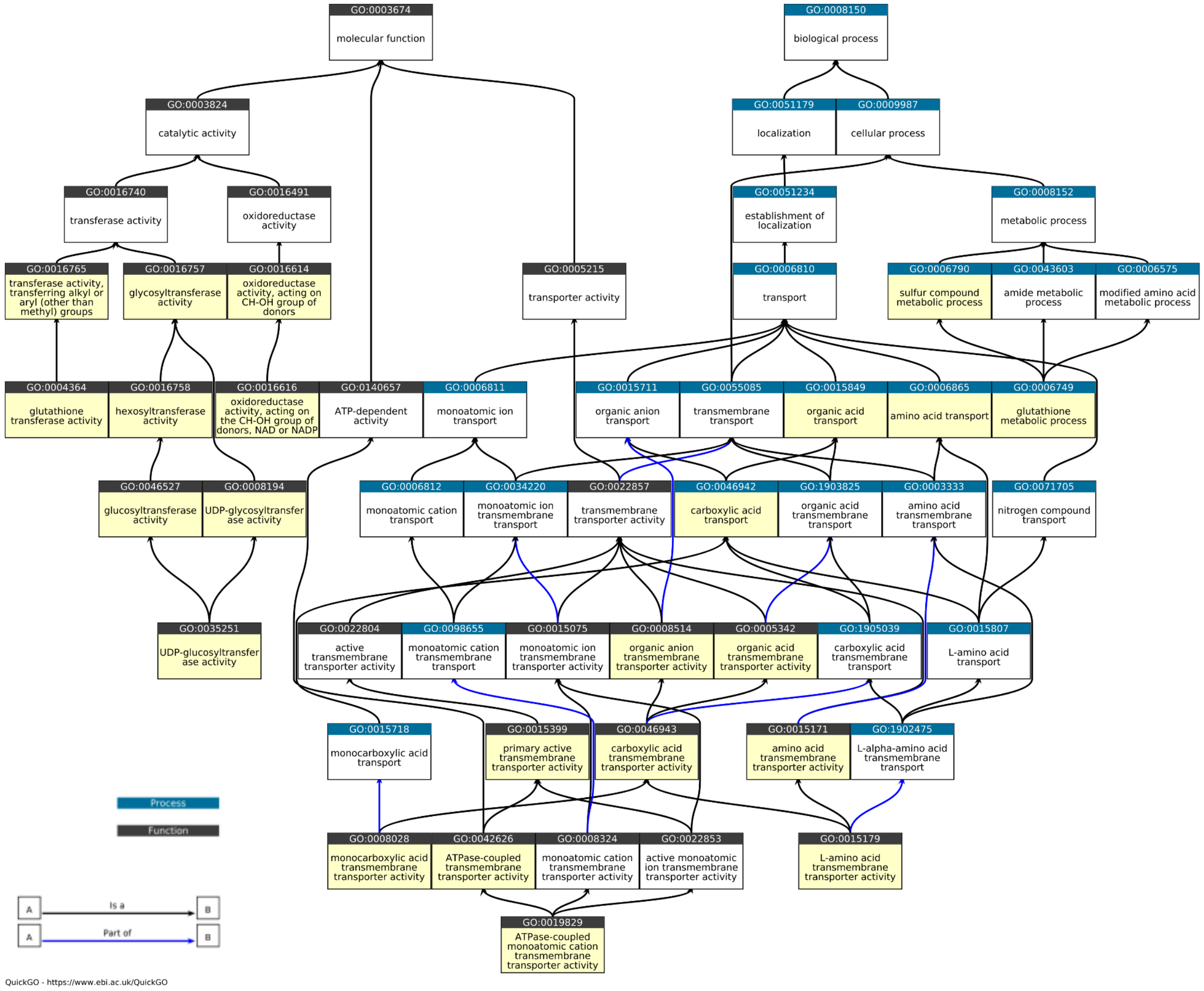

**Figure S4**. Example maximum likelihood phylogenetic trees supporting inferred duplication events. Bootstrap support values for key nodes are shown.

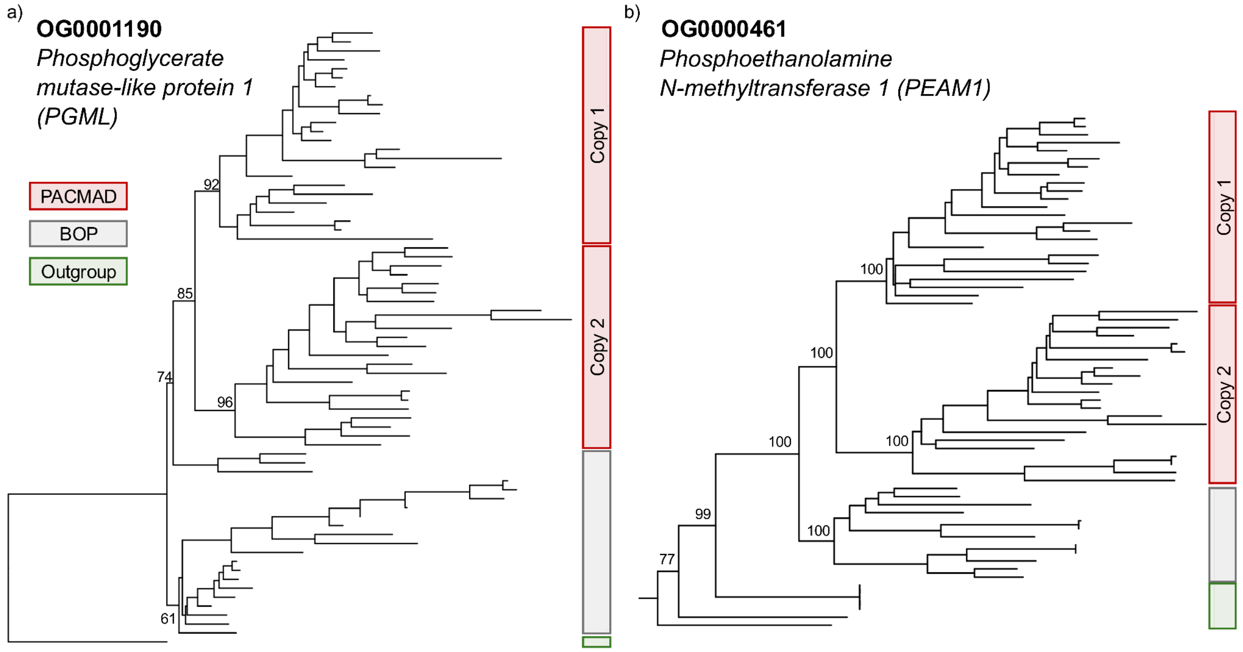

**Figure S5**. Overall gene copy number in the orthogroups that contained the 15 core PACMAD-specific duplication events. Each point represents a different species, and an asterisk indicates orthogroups with significantly higher gene copy number in the PACMAD compared to the BOP clade. Boxplots show the median and interquartile range, whiskers connect the last points within 1.5× the interquartile range.

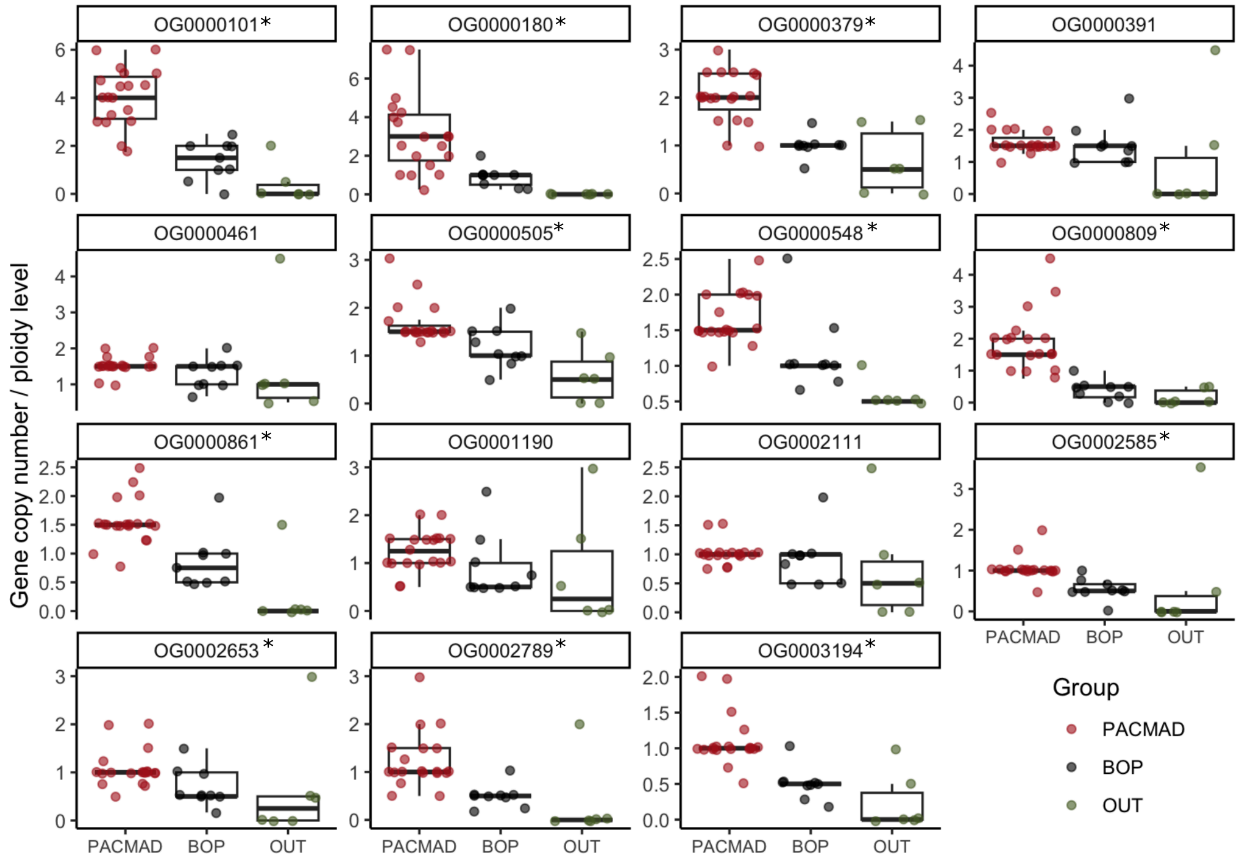

**Dataset S1.** Phylogenetic tree files generated by OrthoFinder
